# Fusion and fission events regulate endosome maturation and viral escape

**DOI:** 10.1101/2020.09.14.295915

**Authors:** Mario Castro, Grant Lythe, Jolanda M. Smit, Carmen Molina-París

**Author notes:** These authors contributed equally to this work.

## Abstract

Many intra-cellular processes rely on transport by endosomes. Recent experimental techniques have provided insights into organelle maturation and its specific role in, for instance, the ability of a virus to escape an endosome and release its genetic material in the cytoplasm. Endosome maturation and dynamics depend on GTPases called Rabs, found on their membrane. Here, we introduce a mathematical framework, combining coagulation and fragmentation of endosomes with two variables internal to each organelle, to model endosomes as intra-cellular compartments characterised by their levels of (active) Rab5 and Rab7. The key element in our framework is the “per-cell endosomal distribution” and its its dynamical equation or Boltzmann equation. The Boltzmann equation, then, allows one to deduce simple equations for the total number of endosomes in a cell, and for the mean and standard deviation of the Rab5 and Rab7 levels. We compare our solutions with experimental data sets of Dengue viral escape from endosomes. The relationship between endosomal Rab levels and pH suggests a mechanism which can account for the observed variability in viral escape times, which in turn regulate the viability of a viral intra-cellular infection.

**Author summary:** Endosomes are intra-cellular receptacle-like organelles, which transport endocytosed cargo upon internalisation from the plasma membrane. These early endosomes, also known as sorting endosomes, mature to late endosomes, with a lower pH than early ones, as a consequence of the intricate dynamics of a family of molecules called Rabs. Viruses exploit this endosomal pH drop to their advantage. Here we bring together experimental data on Dengue viral escape times from endosomes and a novel mathematical framework inspired by the theory of droplet coalescence, to improve our understanding of endosome maturation, and in turn to quantify the large variability of viral escape times. This mathematical framework can easily be generalised to model the dynamics of other intra-cellular organelles, such as mitochondria or the endoplasmic reticulum.

## Introduction

Endosomes are enigmatic organelles which regulate intra-cellular cargo trafficking [1]. These (literally) *inner bodies* are dynamic in movement and decorated [2]. They can merge; that is, they can undergo fusion^1^. Endosomes can also break up (fission) [3] or kiss and run [4], in a random choreography that allows the cell to sort, recycle or degrade internalised molecules. This endosomal maturation programme requires a dramatic transformation of these organelles: from early endosomes, to late ones, recycling ones and degrading lysosomes [1, 5]. In recent years, a lot of effort has been devoted to understand entry and subsequent intra-cellular viral genome release [6]. Current consensus suggests that many viruses, independently of their entry pathway, are delivered to the so-called early endosomes, and thus, enter an intricate network of endosomes: from early to late endosomes, until the endosomal pH is low enough to trigger viral membrane fusion and escape to the cytosol [7, 8]. In this way, endosomes become one of the main vehicles for viral intra-cellular trafficking in the infected cell, bringing viral genomic cargo from the plasma membrane to the cytosol.

For many viruses, escape is preceded by a sudden drop in the endosomal pH [6, 7, 9]. This often occurs in late endosomes, where in fact, high levels of the GTPase Rab7 have been pointed out as responsible for viral escape [10, 11]. This rather interesting feature seems to indicate that by tracking the endosomal Rab decoration, one is in fact, also following the maturation history of individual endosomes. Rab5 is a marker of early endosomes and Rab7 of late ones [12]. Besides their role as experimental markers, different Rab molecules have been identified as central regulators of the endosomal transport pathway [13]. Of particular relevance, is the fact that endosomal pH has been experimentally linked to different levels of Rab5 and Rab7 [5, 13–15]. Given the large variability observed in viral escape times [9, 11, 16], it is timely to ask ourselves the following questions: (i) is pH alone the main trigger of viral escape, (ii) is the pH dependence gradual (analogical) or abrupt (digital), and (iii) can endosome maturation and dynamics explain the observed heterogeneity in the distribution of viral escape times. The first question has been quantitatively addressed, experimentally and theoretically, in recent work [10, 17, 18]. In this paper, we aim to provide answers to the three questions above. We do so by making use of a novel mathematical framework to describe the population dynamics of endosomes containing endocytosed viral particles. The mathematical approach presented here includes the contribution to endosome maturation and dynamics from fusion and fission events, as well as those molecular processes related to Rab recruitment and endosome acidification. In this way, the framework allows one to characterise the mechanistic details of endosomal maturation, yet reduces the mathematical complexity required in order to explain the experimental data.

### The biology of endosomal Rab5/Rab7 dynamics

Early endosomes (Rab5-positive) undergo a progressive replacement of Rab5 with Rab7 [3]; a process that involves several molecules and chemical reactions (see a schematic summary in Fig. 1, adapted from Ref. [19]). Specifically (see Ref. [19] for details), Rab5-positive endosomes (a signature of early endosomes) gradually become Rab7-positive endosomes (namely, late endosomes) [3]. The complete *endosomal maturation* process then replaces the Rab5 decoration with a Rab7 one. At the molecular level (see Fig. 1), Rab5 (in its two conformations, Rab5:GDP and Rab5:GTP, inactive and active, respectively), Rab7, guanine nucleotide exchange factors (GEFs), GTPase-activating proteins (GAPs), and effector molecules are interconnected on the endosomal membrane [20]. The conversion from Rab5 to Rab7 is a consequence of programmed and simultaneous changes in the nucleotide cycle of both Rab5 and Rab7, which shuttle between inactive, GDP-bound, and active, GTP-bound, conformations. GEFs catalyse the exchange of GDP into GTP and GAPs catalyse the hydrolysis of GTP into GDP [21]. Other positive/negative feedback loops are summarised in Fig. 1. We note that for the purposes of this manuscript the molecular reactions included in Fig. 1 will be encoded in the specific choices for the *molecular currents*. Besides their role in the dynamics of Rab molecules, endosomes also interact with other nearby endosomes. In particular, they can undergo fusion and fission [22], or kiss-and-run [4] events. The latter process occurs when Rab5 restricts the complete fusion of endosomes, allowing the exchange of solutes between them, but without complete intermixing of their membranes [4]. Here, we shall only consider endosomal fusion and fission events.

**Fig 1.**
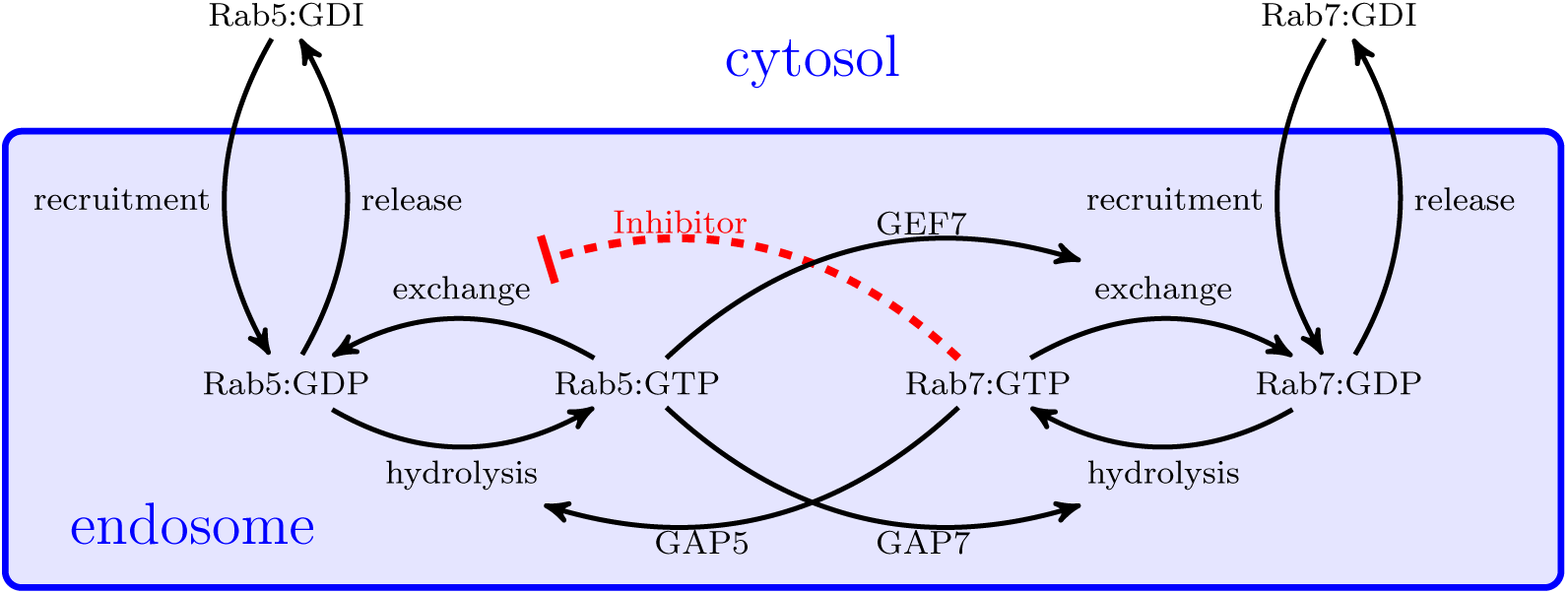
Schematic summary of the reactions which decorate the endosomal membrane. They regulate the levels of Rab5/Rab7 in the cytosol and at the membrane in inactive (Rab-GDP) and active forms (Rab-GTP). GDI means GDP-dissociation inhibitor (see Ref. [19] for details). Mathematically, Eq. (10) and Eq. (11), capture the main molecular interactions. For instance, the red dashed line is modelled by the term proportional to *v*_57_ *x*_7_.

### Mathematical framework of endosome dynamics and maturation

Mathematical models have helped identify the main mechanisms involved in the endosomal maturation process. For instance, some regulatory proteins, *e*.*g*., SNAREs, Rabs and other coating proteins, have been linked to the formation, degradation and renewal of endosomes. There is some consensus on what mechanisms are involved in those processes, although their individual quantitative contribution seems to be elusive [15, 23]. In this section, we summarise the biology of endosomal Rab dynamics, as well as previous mathematical modelling efforts.

Most mathematical models of endosome dynamics (with the notable exception of the model in Ref. [24] that pioneered the approach developed here) make use of a set of ordinary differential equations (ODEs) to describe the incorporation of different Rab molecules to the endosome. This poses a practical problem since, in all cases, the data used to parameterise the mathematical models is averaged over all the endosomes. However, as we mentioned above, fusion and fission events couple two nearby endosomes and, hence, endosomal averages might not be appropriate to describe the pH of an individual endosome. To illustrate why traditional ODE models do not capture this variability properly, let us consider an extreme case with two endosomes with pH 4 and 7, respectively, and let us assume that viral escape occurs for pH below 5. The average endosome pH will be 5.5 in this case, so escape will not occur, which contradicts the fact that it will in one of the endosomes, but not the other. Hence, endosomal averaging can smooth out fast changes and lead to the conclusion that escape is a continuous process rather than mediated by a pH threshold [23]. At the other end of the modelling spectrum, the biochemistry of proton exchange has been used to describe the thermodynamics of endosome pH acidification [25]. Also, the authors of Ref. [14] consider the acidification of individual endosomes, without considering interactions between them. Finally, it is worth mentioning that, in Ref. [26] the authors made use of a hybrid method, which includes differential equations for active Rab5, inactive Rab5 and the total number of endosomes. Specifically, the model considers the mean number of Rab5 molecules per endosome and assumes that Rab5 exchange and activation between cytosol and endosomes is proportional to the number of endosomes (see Eqs. (75)-(78) in Section 3 of the Supporting information, which were proposed by the authors of Ref. [26]). Note that Eq. (78) incorporates the role of fusion and fission events in terms of the mean number of endosomes. Thus, this *mean field* approach is unable to describe individual endosome dynamics. While this can be a valuable approach to understand the organelle network, it does not capture endosome heterogeneity, which is essential if we want to understand and describe the observed experimental variability. A suitable approach to resolve this issue would be the use of agent-based models (ABM) [27], in which each endosome can be simulated, as an agent, and is described in terms of its molecular content (or cargo). However, ABMs are often computationally expensive or not amenable to parameter inference.

Our approach here takes a different route beyond deterministic descriptions of endosomal dynamics at the cellular level [19, 26, 28, 29], or at the single endosome level [14, 15, 23]. As we have discussed earlier, the former presumes that Rab dynamics at the cellular scale can be studied from the observation of endosomal averages, and, the latter, focuses on biochemical reactions on the endosome membrane, which do not include the interaction between endosomes. To overcome these limitations, we introduce a model inspired by the dynamics of droplet coalescence, also known in the physics literature as the Smoluchowski equation [24, 30, 31]. Our approach allows one to recover previous phenomenological models [19, 26] under certain mathematical assumptions (see Section 3 of the Supplementary information). Mathematically, these models are based on an integro-differential equation which can describe cells [32–34] or endosomes [24, 35]. A difficulty with phenomenological models is their lack of mathematical tractability. In order to address this issue, here, we make use of a formalism that enables one to describe the dynamics of the number of endosomes and their individual cargo of (active) Rab5 and Rab7 molecules. Furthermore, by means of the two-dimensional Laplace transform, the formalism naturally leads to analytical expressions for the rate equations (ODEs) of interest. Once the rate equations have been derived, and together with experimental data [8], we test the validity of these equations and discriminate between two different mechanistic hypotheses [36]. Equipped with this mathematical framework, we are able to simplify some of the underlying assumptions used in other models (see Section 3 on the Supplementary information) and perform a model selection analysis (making use of experimental data). Our interest is to directly emphasise the underlying molecular mechanisms rather than certain mathematical choices; this can be done since our framework avoids *ad hoc* reaction terms based on Hill or logistic functions (see the Results section).

Finally, and going back to our initial discussion on endosome acidification, we note that the quantification of intra-cellular pH can be performed with the use of DNA nanomachines [37] or by luminescence markers of acidity acting as cargo transported by the endosome [38]. Since the experimental error can be as large as ΔpH ∼ 1, these experiments might not be accurate enough in the case of abrupt pH changes. To overcome these experimental limitations, several approaches have been proposed to understand the interplay between Rab decorations and endosomal pH [14, 23]. These approaches, in a nutshell, provide a mechanistic model of proton dynamics and an interpolation model between early and late endosomes, by means of an equation of the following kind:

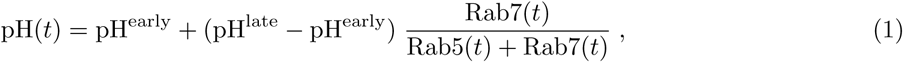

where Rab5(*t*) and Rab7(*t*) have to be understood at the single endosome level, not at the single cell level. We shall make use of this equation to derive the time course of endosomal pH, pH(*t*), by providing the dynamics of Rab5(*t*) and Rab7(*t*) as derived from our general mathematical framework.

### Mathematical modelling hypotheses

We take into account the experimental evidence discussed in the preceding sections and now describe the biological mechanisms that will be included in our mathematical formalism.

1. **Endocytosis (endosome generation)**: new endosomes are formed by the invagination of the cell membrane. Those newly created endosomes (containing endocytosed viral particles) do not have any Rab5 or Rab7 molecules. Progressively, the endosomes become *decorated* with Rab5 and, subsequently, with Rab7.
2. **Endosome degradation/removal**: endosomes can fuse with a lysosome and thus, be removed from the cytosol. In our case, there exists a second form of endosome removal due to viral escape.
3. **Fusion (coalescence)**: fusion of two endosomes involves the merging of their membranes and, as a consequence, the Rab5 and Rab7 molecules are shared [3] (see Fig. 2 for a schematic picture of this process).
4. **Fission**: endosomes can divide, thus splitting the amount of Rab5 and Rab7 between the two newly created endosomes. When fission occurs, the total amount of Rab5 and Rab7 is shared randomly between the two newly created endosomes (see Fig. 2 for a schematic picture of this process).
5. **Rab5/Rab7 activation/deactivation**: Rab molecules on the endosomal membrane can be activated after prior incorporation of inactive Rab:GDI from the cytosol. A schematic summary of the reactions of Rab activation/deactivation can be found in Fig. 1.

**Fig 2.**
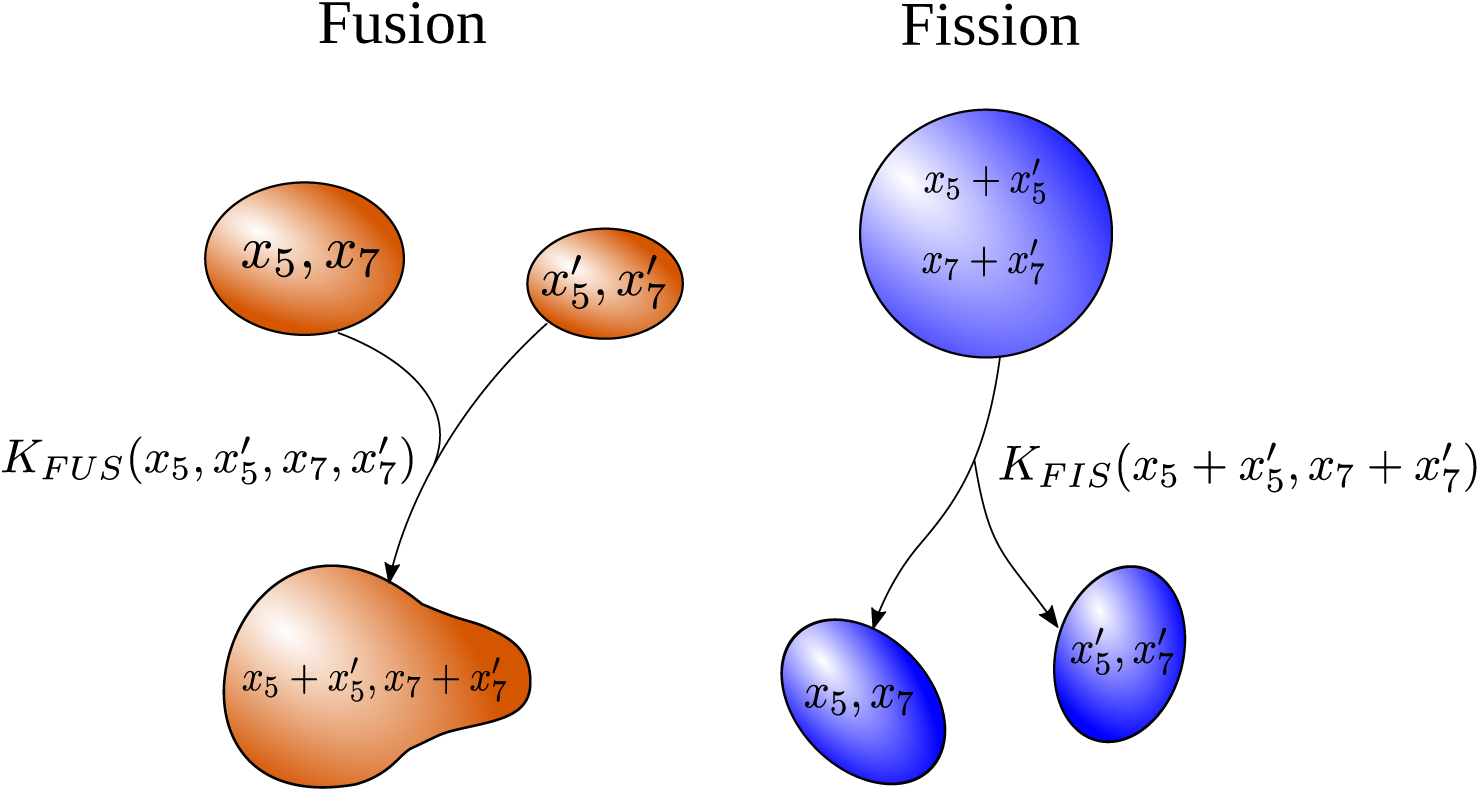
Left: two endosomes with different levels of active Rab5 and Rab7, (*x*_5_, *x*_7_) and 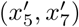, respectively, merge into a new endosome with levels of active Rab5 and Rab7, 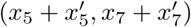, with rate 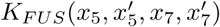. Right: an endosome with levels of active Rab5 and Rab7 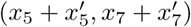 splits into two new endosomes of levels (*x*_5_, *x*_7_) and 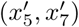, respectively, with rate 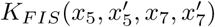.

### Mathematical model of endosomal maturation and dynamics

In this section, we provide a mathematical description of the biological mechanisms discussed in the previous section. We consider a single cell containing a collection of endosomes. Each endosome is characterised by its active Rab cargo, technically, [Rab5:GTP] and [Rab7:GTP], respectively, (*x*_5_, *x*_7_) at time *t*. We assume *x*_5_ and *x*_7_ are real numbers (greater or equal to zero) and introduce the “per-cell endosomal distribution”, *n*(*x*_5_, *x*_7_; *t*), which is a function of *x*_5_, *x*_7_ and time, *t*. In fact, the mean number of endosomes at time *t* with Rab cargo (*x*_5_, *x*_7_), such that *a*_5_ *< x*_5_ *< b*_5_ and *a*_7_ *< x*_7_ *< b*_7_ is given by

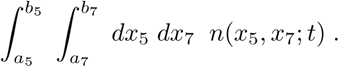

Without loss of generality and, for the sake of simplicity, we make use of the number of Rab molecules per endosome instead of the number density or the molar concentration. We now consider the contribution of each of the five processes just introduced to the time evolution (or dynamical equation) of *n*(*x*_5_, *x*_7_; *t*).

1. **Endocytosis (endosome generation)**: newly created endosomes have zero levels of active Rab5 and Rab7. Mathematically, endosome generation is described by a source term of the form

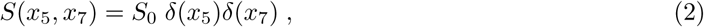

where *S*_0_ is a real positive constant with dimensions of number of endosomes per unit time and the symbol *δ*(·) stands for the Dirac delta function.
2. **Endosome degradation/removal**: considering the two possible causes of endosome removal (fusion with a lysosome or viral escape out of the endosome), we assume that the endosomes can be removed at any stage of the maturation process. Hence, we assume a constant removal (or *death*) rate proportional to the number of endosomes, that is independent of the Rab cargo. We write

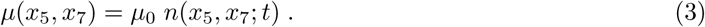
3. **Fusion (coalescence)**: when two endosomes with Rab cargo given by (*x*_5_, *x*_7_) and 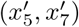, respectively, fuse, they share their cargo, so the total number of each Rab cargo of the newly formed endosome will be 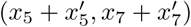. The contribution to the time derivative of *n*(*x*_5_, *x*_7_; *t*) is

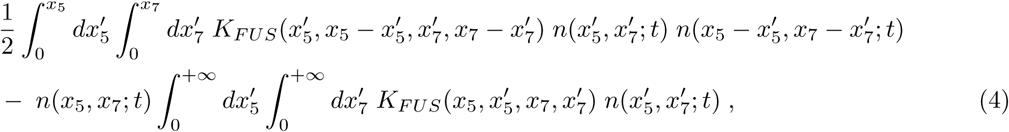

where the first term represents the net *gain* of endosomes with Rab levels (*x*_5_, *x*_7_) and the second one represents the *loss* of endosomes with total levels (*x*_5_, *x*_7_) after fusion with other endosomes. The function 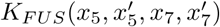 is referred to as the fusion kernel. It dictates the rate of endosomal fusion, and it clearly depends on the endosomal levels of active Rab molecules. A central feature of our model is the consideration of both fusion and fission events. Fusion is enhanced in early endosomes so the rate of fusion correlates positively with the levels of Rab5 and negatively with those of Rab7 [3]. For simplicity, we assume that the fusion rate is a linearly increasing function of Rab5 and a linearly decreasing function of Rab7. Hence, we propose

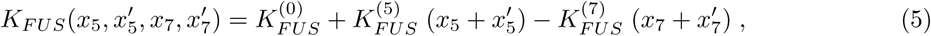

where 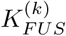 are constants for *k* = 0, 5, 7. As shown in Ref. [39], endosomes need to be spatially close in order to merge. As we are not modelling the intra-cellular endosome spatial location explicitly, the latter equation favours fusion of early endosomes (higher *x*_5_ increases the overall fusion rate) and reduces fusion in late endosomes (it decreases for higher *x*_7_). Yet our model does not preclude fusion events between early and late endosomes [8].
4. **Fission:** similarly to fusion, fission can be described introducing a kernel function, as follows

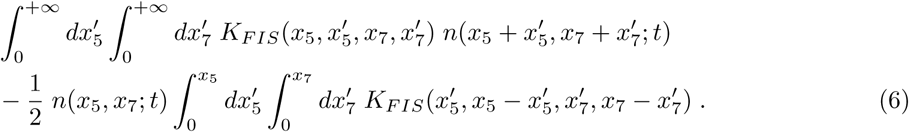

The first term is the *gain* due to the fission of a larger endosome leading to two endosomes, one of them with Rab levels (*x*_5_, *x*_7_). The second one is a *loss* term due to the fission of an endosome with Rab levels (*x*_5_, *x*_7_). Endosomal fission is less well understood that fusion. In Ref. [3] it is suggested that fission occurs randomly at any stage of maturation. Thus, we consider that fission is independent of the number of Rab5 or Rab7 molecules, but that it is not necessarily symmetric (namely, when an endosome splits, the amount of Rab going to each daughter endosome can be different). Mathematically, we propose

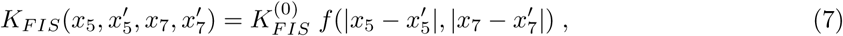

where, by the symmetric properties of the fission *kernel* [30], the function *f* satisfies the normalisation condition, *f* (0, 0) = 1 and is symmetric in its arguments. The simplest case one can consider is symmetric fission; namely, we write

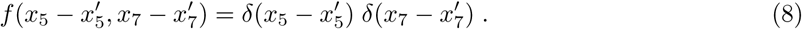

This choice for *f* means that 50% of each cargo is equally shared between daughter endosomes. The contribution to the time derivative of *n*(*x*_5_, *x*_7_; *t*) is then

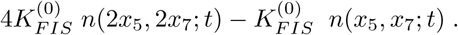
5. **Rab5/Rab7 activation/deactivation in an endosome**: in Ref. [19], the authors considered two competing hypotheses for Rab5/Rab7 activation/deactivation (see Fig. 1). The first one is the *toggle-switch* model and consists in a weakened repression of Rab7 by Rab5, described by a logistic term. In the second one, the *cut-off switch* model, Rab7 activation strongly suppresses Rab5. In order to identify which hypothesis was more compatible with the experimental data, the authors introduced a modular model where a certain mechanisms could be explained making use of different mathematical functions (see Supplementary information 1 of Ref. [19]). The drawback of this type of exhaustive model comparison is that specific fitting algorithms had to be adapted to infer model parameters from the data [28]. We note that the total number of endosomes does not change with the activation/deactivation of Rab molecules. This fact can be naturally expressed in terms of a conserved quantity. We also note that we model the number of Rab molecules in a given endosome rather than its concentration, since the latter one might be affected by fusion and fission events, where the volume or the area of the endosome can significantly change. In the absence of other mechanisms, this conservation law can be expressed in terms of a *molecular current*. The contribution of the dynamics of *x*_5_ and *x*_7_ inside each endsosome to the time derivative of *n*(*x*_5_, *x*_7_; *t*) is equal to minus the divergence of a current 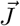:

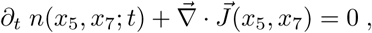

where

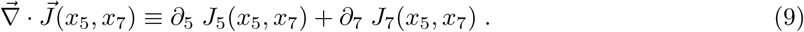

In the previous equation we have introduced the notation ∂_*k*_ to indicate 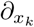, with *k* = 5, 7. We have also made use of the notation 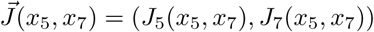. One must supplement Eq. (9) with *constitutive equations* for the currents, *J*_5_(*x*_5_, *x*_7_) and *J*_7_(*x*_5_, *x*_7_). In this paper we are going to assume that the currents are proportional to the total number of endosomes, *n*(*x*_5_, *x*_7_; *t*). This implies that *J*_5_(*x*_5_, *x*_7_) = *v*_5_(*x*_5_, *x*_7_) *n*(*x*_5_, *x*_7_; *t*) and *J*_7_(*x*_5_, *x*_7_) = *v*_7_(*x*_5_, *x*_7_) *n*(*x*_5_, *x*_7_; *t*), where we have introduced the velocities *v*_5,7_(*x*_5_, *x*_7_). The *velocities v*_5,7_ are generic functions of *x*_5_ and *x*_7_ that need to be prescribed according to the underlying biology of Rab5/Rab7 activation/deactivation discussed earlier. Following Refs. [19, 26], we assume that both molecules evolve and are coupled to each other (see Fig. 1). In other words, the concrete form of the functions *v*_5,7_(*x*_5_, *x*_7_) will be determined by performing model selection and thus, identifying the underlying biological mechanisms. Mathematically, Rab5 and Rab7 interact via positive/negative feedback loops. We include these molecular interactions in the current, 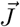, and assume a linear dependence on the number of Rab molecules. For the *cut-off switch* model one has [19]

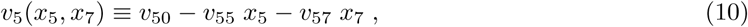

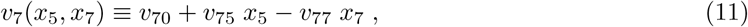

where the choice of the signs in the coefficients *v*_*ij*_ is determined by the network of interactions in Fig. 1. For instance, the inhibition described by the red dashed arrow is captured by the term −*v*_57_ *x*_7_. For the *toggle-switch* model one has *v*_57_ = 0 [19]. Namely, levels of Rab7 do not affect levels of Rab5, and thus, the velocity *v*_5_(*x*_5_, *x*_7_) does not depend on *x*_7_. Yet, the model is non-linear, since it includes two logistic terms. The non-linear terms encode inhibitory mechanisms, as for example, the terms proportional to *v*_55_ and *v*_75_. For the *toggle-switch* model one has [19]

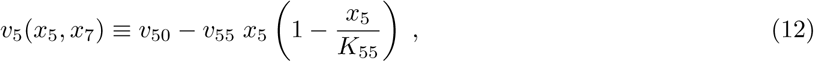

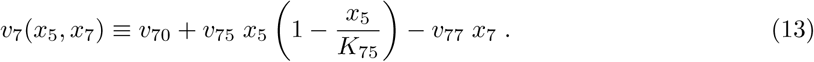

The parameters *K*_55_ and *K*_75_ are *carrying capacities* that encapsulate the inhibitory behaviour of Rab5 in the toggle-switch model. As a consequence, we shall show (see Eq. (26)), that the ODE for the endosomal average of Rab5 does not contain the inhibitory feedback proportional to *v*_57_ present in Eq. (23).

### Boltzmann equation for the endosomal distribution

We now combine the mathematical considerations described in the previous section to derive a dynamical equation for *n*(*x*_5_, *x*_7_; *t*). Before we do so, we need to discuss the relationship between the creation of endosomes and the current 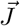. As newly created endosomes have zero levels of Rab5 and Rab7 (see Eq. (2)), the following boundary conditions for the current 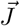 must be fulfilled:

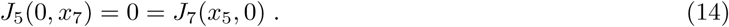

Intuitively, these equations mean that the Rab5-associated current of endosomes with non-zero levels of Rab7 (first equation) and the Rab7-associated current of endosomes with non-zero levels of Rab5 (second equation) have to be zero.

We can now write the evolution equation for *n*(*x*_5_, *x*_7_; *t*). We have

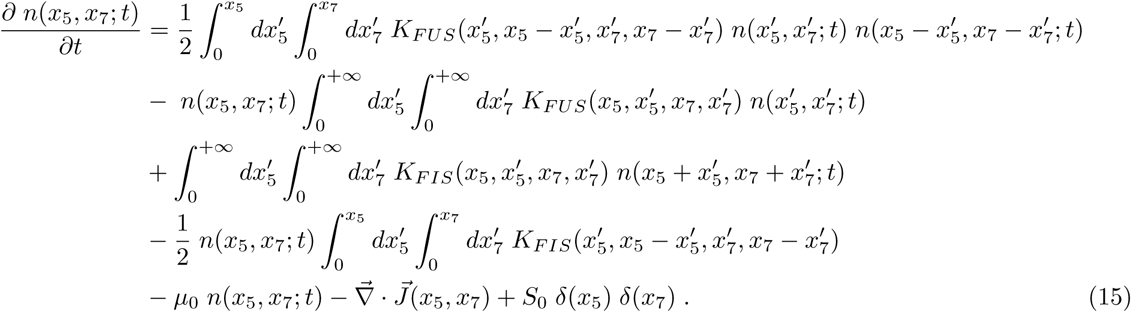

We refer to this equation as the Boltzmann equation for the endosomal distribution *n*(*x*_5_, *x*_7_; *t*). Although this precise equation has not been proposed before, simplified versions with fewer biological mechanisms [24] or in fewer dimensions [34, 40] have been studied in different contexts.

### Equations for the moments of the Boltmann distribution

Equation (15) is a non-linear integro-differential equation that is, in principle, analytically intractable. However, it can be simplified under some assumptions that should be carefully scrutinised together with experimental data. In practice, in many experimental conditions only the time evolution of the mean number of molecules of different species (including the total number of endosomes) is attainable. To this end, it is convenient to introduce the first order *moments* of the distribution *n*(*x*_5_, *x*_7_; *t*). In particular, we introduce

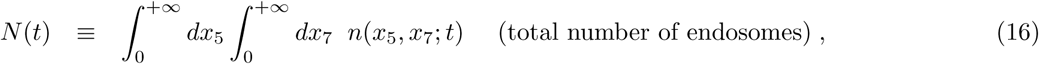

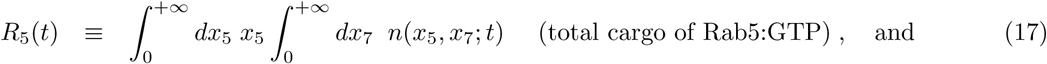

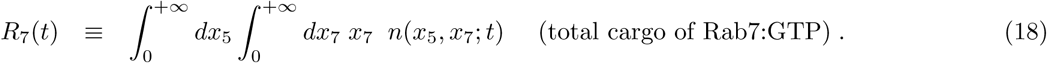

We can also define second order moments of the distribution as follows

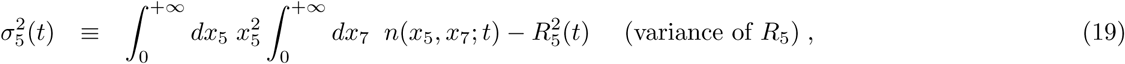

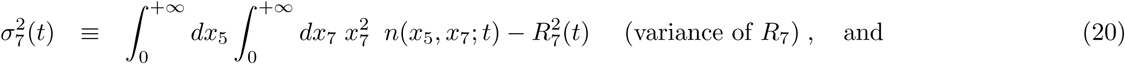

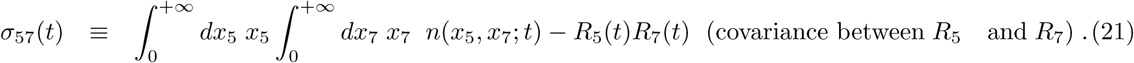

If we make use of the two-dimensional Laplace transform (for details, see Section 1 in the Supporting information), we obtain an ODE for the first order moments defined above. For the *cut-off switch* model one can show

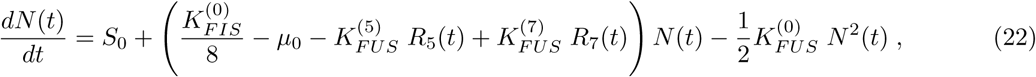

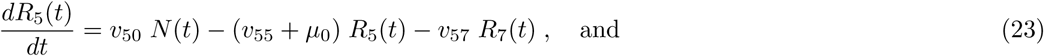

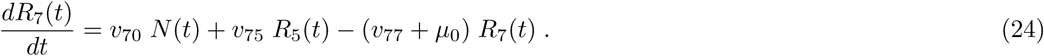

For the *toggle-switch* model the ODEs for the first moments can be shown to be

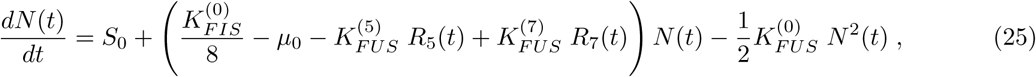

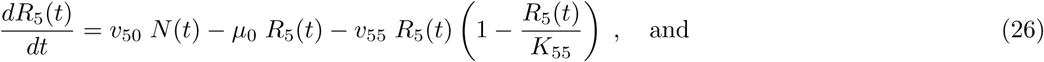

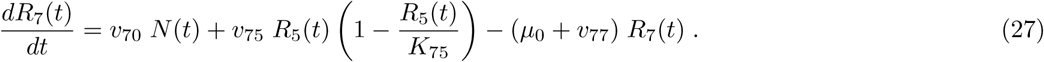

We note that as expected, Eq. (25) and Eq. (22) are the same in both models. We also note that Eq. (15) allows one to derive the dynamical non-linear equation of Refs. [19, 26] (for further details, please, see Section 3 in the Supporting information).

## Experimental data

We make use of experimental data obtained by labelling Dengue viral particles (DENV) with the lipophilic fluorescent probe DiD, as previously reported in Ref. [8] and reproduced in Fig. 3. We use the normalised number of *probes* (DENV viral particles) colocalised with endosomal Rab5 and Rab7 as an estimate of the number of Rab molecules in endosomes with endocytosed DENV. Here, we are not interested in the DENV lifecycle, but the viral particles will serve as markers to track endosomal dynamics, since in the experiments viral particles were colocalised with fluorescent markers for endosomal Rab5 and Rab7 molecules. Overall, 51 escape events were analysed in order to quantify the levels of Rab5 and Rab7 (see Fig. 3). Analysis of those 51 cases revealed that, in spite of the fact that most escape events took place in early endosomes (86%), a non-negligible number of events, 14%, took place in Rab5/Rab7-positive intermediate endosomes [8]. In addition, tracking fluorescently labelled endosomes allowed the authors to show that almost half of the endosomes skipped several steps of the maturation process by merging with existing Rab7-positive endosomes (precisely, 45%) [8]. Finally, 30% of the tracked endosomes underwent fission events at different stages of their trajectory in the cytoplasm. This supports our choice for a constant fission rate, termed *splitting* in Ref. [8], which does not depend on the stage of maturation of the endosome, as defined by its Rab cargo (*x*_5_, *x*_7_).

**Fig 3.**
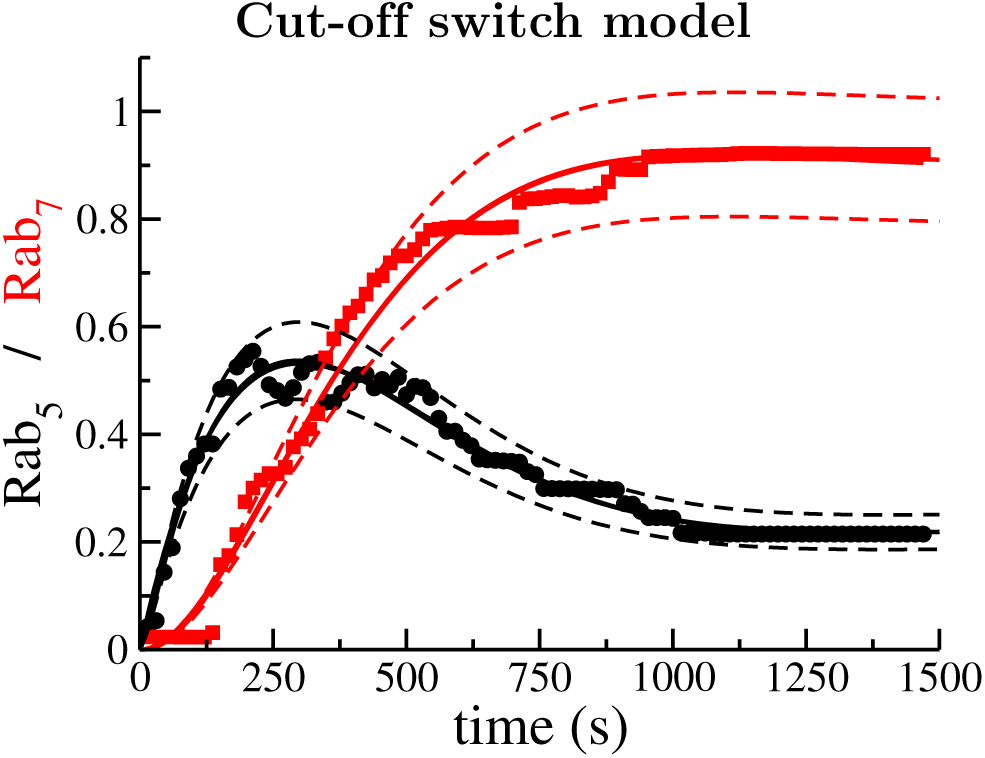
Comparison of model predictions (solid lines) as described by Eqs. (22)-(24) and the experimental data (black circles and red squares) from Ref. [8]. Solid lines correspond to the normalised number of endosomal Rab5 and Rab7 molecules. Dashed lines correspond to the mean *±* one standard deviation, as derived from Eqs. (51)-(53).

## Results

### Data and mathematical modelling support the cut-off switch hypothesis

Fig. 3 shows the experimental data (black circles and red squares) from Ref. [8] and the result of fitting the data to the mathematical model (cut-off switch) described by Eqs. (22)-(24) (solid lines). A sensitivity analysis of the cut-off switch model (see Table 2 of Section 2 in Supporting information) reveals that the most sensitive parameters are, in order of relevance, 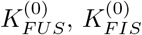, *v*_50_ and *v*_57_, and the least relevant, 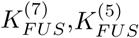, *µ*_0_ and *S*_0_. The results from our sensitivity analysis is rather interesting since it shows that fusion and fission are essential to understand the experimental data, but that the corrections to the constant term of the fusion kernel, 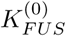, which are proportional to 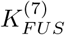 and 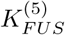 and depend on the Rab cargo, are negligible. Thus, it seems that a simpler model than the cut-off switch can be used to explain the data, as we show in Section 2 in Supporting information (see also A quasi-linear approximation to describe experimental data section below and Fig. 4C). Furthermore, the source term of new endosomes with zero levels of Rab5 and Rab7 does not significantly affect the dynamics of endosomal Rab5 or Rab7. Numerical integration of Eq. (22) shows that, independently of the initial condition, *N* (0), the number of endosomes quickly approaches a steady state (see Fig. 6). Taking into account the order of magnitude of the parameters in Table 1 and the results of the sensitivity analysis, we find that

**Table 1.** Best-fit parameters obtained by three different methods as implemented in the software Copasi [47] for: i) the cut-off switch model defined by Eqs. (22)-(24), ii) the reduced model defined by Eq. (69) and Eq. (70), and iii) the toggle-switch model degined by Eqs. (25)-(27). All parameter values are given in units of seconds^−1^.

| Parameter | Cut-off | Reduced | Toggle-switch |
| --- | --- | --- | --- |
| $K_{FIS}^{(0)}$ | 4.54 | 4.48 | 0.7723 |
| $K_{FUS}^{(0)}$ | 0.0032 | 0.0032 | 0.0039 |
| $v_{50}$ | $1.3 \times 10^{-5}$ | $1.2 \times 10^{-5}$ | $1.3 \times 10^{-5}$ |
| $v_{50} N_{ss}$ | 0.0050 | 0.0045 | 0.0050 |
| $v_{57}$ | 0.0040 | 0.0036 | — |
| $v_{77}$ | 0.0013 | 0.0010 | 0.0013 |
| $v_{75}$ | 0.0032 | 0.0039 | 0.00314 |
| $v_{55}$ | 0.0062 | 0.0058 | 0.0062 |
| $v_{70}$ | $1.2 \times 10^{-6}$ | — | $1.2 \times 10^{-6}$ |
| $S_0$ | 0.4389 | — | 0.1586 |
| $\mu_0$ | $1.0 \times 10^{-6}$ | — | $1.0 \times 10^{-6}$ |
| $K_{FUS}^{(5)}$ | $2.0 \times 10^{-6}$ | — | $1.8 \times 10^{-6}$ |
| $K_{FUS}^{(7)}$ | $1.2 \times 10^{-5}$ | — | $1.2 \times 10^{-6}$ |
| $K_{55}$ | — | — | 0.45 |
| $K_{55}$ | — | — | 0.53 |

**Table 2.** Sensitivity analysis of the mathematical model described by Eqs. 22-(24), corresponding to the cut-off switch hypothesis in Fig. 1, as computed by three different methods implemented in the software Copasi [47]. The column *Aggregated* is defined in Eq. 68. The column *Normalised* is the value in *Aggregated* divided by the maximum value in that column. The thin dotted lines are a guide to the eye to separate the most sensitive parameters (top part of the table) and the least (bottom part).

| Parameter | $\Sigma_N$ | $\Sigma_{R_5}$ | $\Sigma_{R_7}$ | Aggregated | Normalised |
| --- | --- | --- | --- | --- | --- |
| $K_{FUS}^{(0)}$ | -0.9986 | -0.9984 | -0.9984 | 1.7294 | 1.0000 |
| $K_{FIS}^{(0)}$ | 0.9990 | 0.9955 | 1.0007 | 1.7293 | 0.9999 |
| $v_{50}$ | 0.0000 | 1.4462 | 0.8411 | 1.6731 | 0.9674 |
| $v_{57}$ | 0.0000 | -1.1510 | -0.6141 | 1.3046 | 0.7543 |
| $v_{77}$ | 0.0000 | 1.0718 | -0.3886 | 1.1401 | 0.6593 |
| $v_{75}$ | 0.0000 | -0.7050 | 0.2265 | 0.7405 | 0.4282 |
| $v_{55}$ | 0.0000 | -0.4462 | 0.1589 | 0.4737 | 0.2739 |
| $v_{70}$ | 0.0000 | -0.2259 | -0.2519 | 0.3384 | 0.1957 |
| $S_0$ | 0.0005 | 0.0005 | 0.0005 | 0.0009 | 0.0005 |
| $\mu_0$ | -0.0001 | 0.0007 | -0.0003 | 0.0008 | 0.0004 |
| $K_{FUS}^{(5)}$ | $< 10^{-4}$ | $< 10^{-4}$ | $< 10^{-4}$ | $< 10^{-4}$ | $< 10^{-4}$ |
| $K_{FUS}^{(7)}$ | $< 10^{-4}$ | $< 10^{-4}$ | $< 10^{-4}$ | $< 10^{-4}$ | $< 10^{-4}$ |

**Fig 4.**
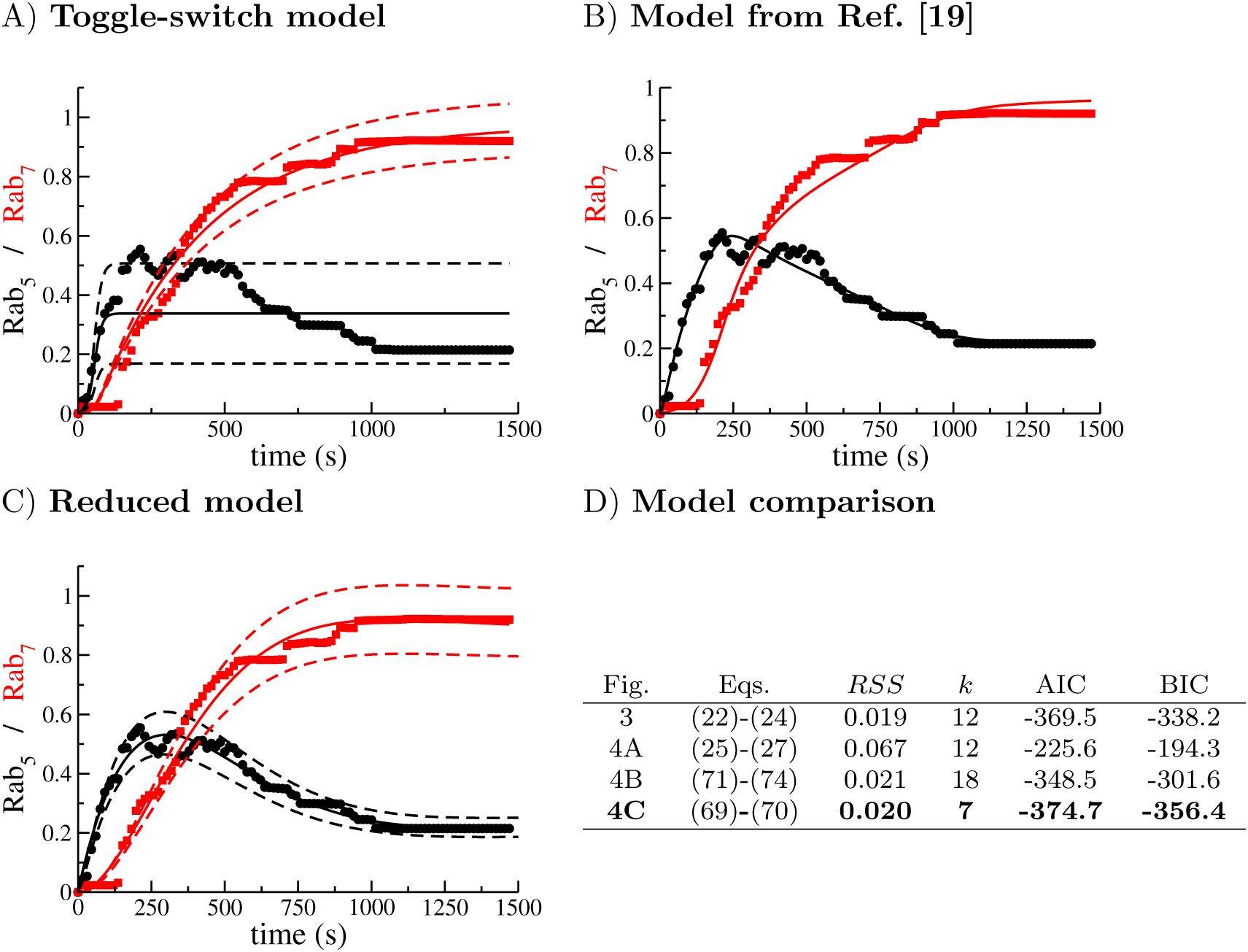
Comparison of different models (solid lines) with the experiments of Ref. [8]. A) Toggle-switch model, Eqs. (25)-(27). B) Model from Ref. [19] (see also Eqs. (71)-(74) in Supporting information). C) Comparison of the sensitivity-based reduced model, Eqs. (69)-(70) (see Section 2 in Supporting information for details). D) Model comparison based on the Akaike Information Criterion, Eq. (29) and the Bayesian Information Criterion, Eq. (30). The selected model according to the minimum AIC and BIC is the one shown in boldface. Dashed lines represent the mean the standard deviation, as described by Eqs. (48)-(49). We note that the formalism introduced here allows one to predict the variance of the estimated solution (panels A and C), unlike traditional ODE modelling approaches, where only the mean can be explained (panel B).

**Fig 5.**
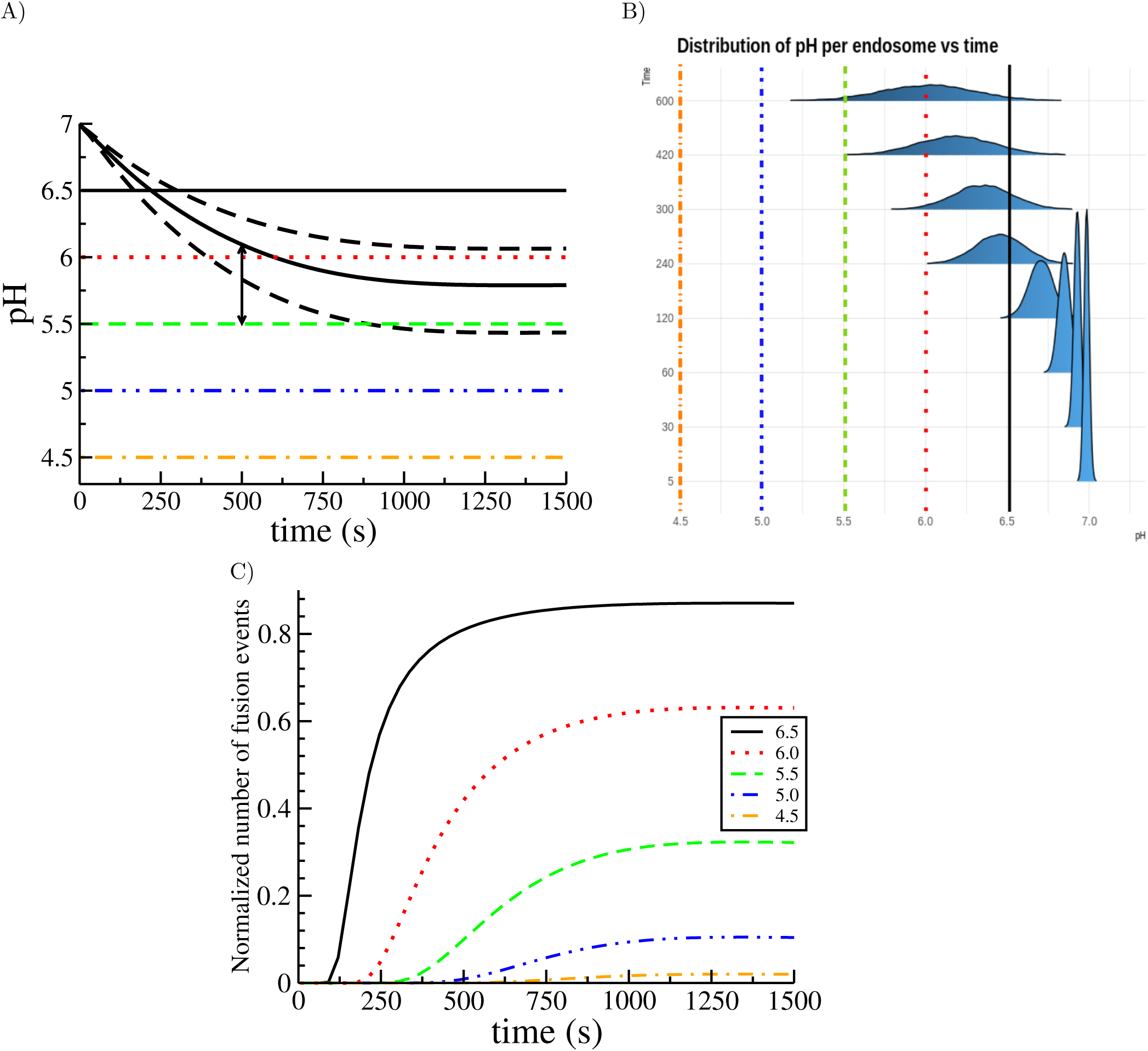
A) Time evolution of the mean endosomal pH (thick curved line) computed from Eqs. (22)-(24) and Eq. (1), and mean *±* standard deviation (computed from Eqs. (51)-(53), curved dashed lines). The horizontal colour lines correspond to different pH thresholds that could be linked to a virus endosomal escape. B) Similar to panel A) but showing the sampled distribution of pH using Eq. (31). Note how the distribution broadens with time, thus increasing the probability of crossing a given pH threshold. C) Normalised number of viral escape events quantified as the probability of pH being below a threshold (see colour-coded legend), making use of Eq. (31), namely, 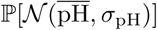 *<*threshold.

**Fig 6.**
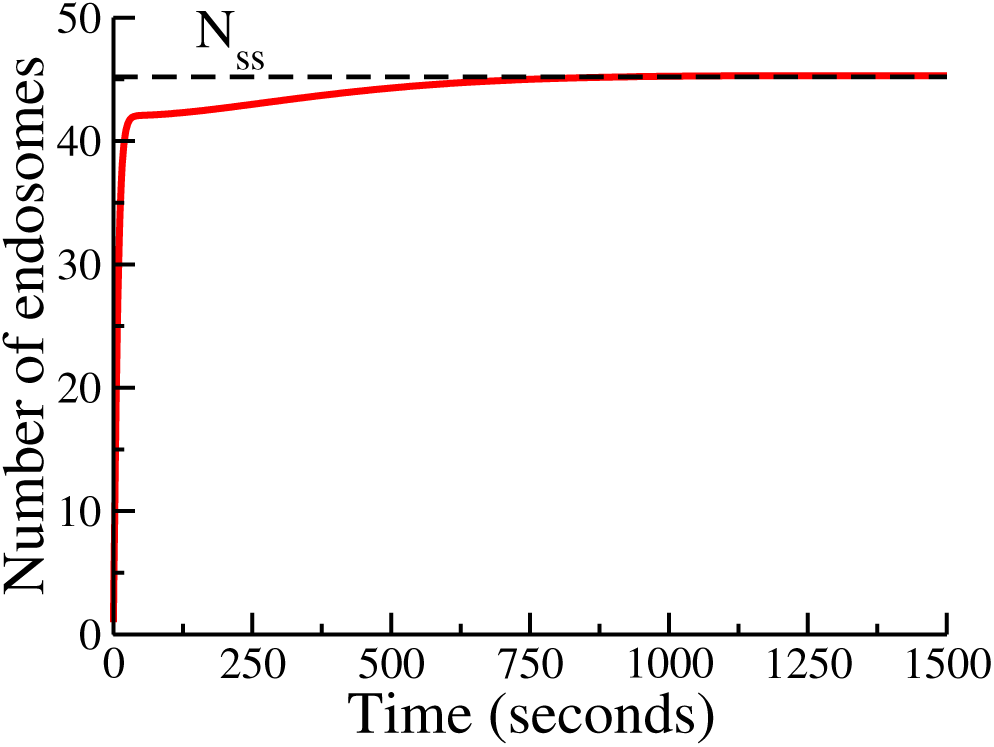
Time course of the number of endosomes, *N* (*t*), for the cut-off switch model (solid red line) and its steady state value, *N*_ss_, (black dashed lines). Note that *N* (*t*) grows rapidly in the first 40 seconds and reaches the steady state value of the reduced model after 500 seconds. The numerical value is extremely close to the value given by Eq. (28).

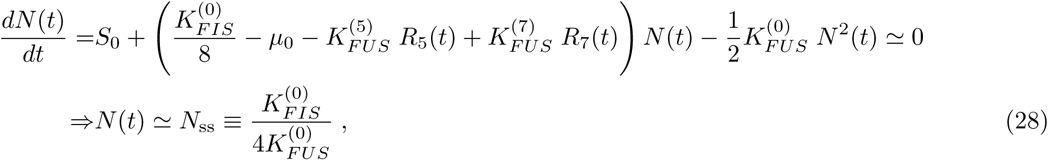

where we have neglected 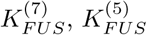, *µ*_0_ and *S*_0_ and assumed *N*_ss_ ≠ 0. As a consequence of *N* (*t*) being almost stationary, the terms *v*_50_*N* and *v*_70_*N* in Eq. (23) and Eq. (24), respectively, are also almost constant. Our theoretical analysis is consistent with our numerical results: Fig. 6 shows that the number of endosomes, *N* (*t*), reaches a value close to its steady state after 40 seconds. Given the best-fit parameters from Table 1, we conclude that fusion and fission events are more relevant for the dynamics of the Rabs than the rate of endosome generation, since we have 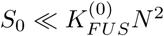 and 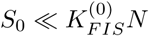. Finally, we now explore the role of the parameter *v*_57_ in the decrease of *R*_5_(*t*) at the time of increase of *R*_7_(*t*). This coefficient encodes the inhibitory effect of Rab7 on the dynamics of Rab5. On the one hand, the sensitivity analysis of the cut-off switch model (see Table 2) gives this parameter a normalised value of 0.7543 and on the other hand, the best-fit parameters from Table 1, suggest a value of 4 *×* 10^−3^ s^−1^ for *v*_57_. These two results together show the qualitative and quantitative importance of this parameter in the cut-off switch model, which in turn, and in light of the experimental data, provide most support to the cut-off switch hypothesis ^2^, in agreement with Ref. [19]. To further test this conclusion, we have also fitted the toggle-switch model, Eqs. (25)-(27), to the experimental data. The results are shown in Fig. 4A. It can be concluded that the toggle-switch model cannot explain the decrease of Rab5 observed in the data. We first note that in Eq. (26) *R*_5_(*t*) does not depend on *R*_7_(*t*). We then argue that for this model, only fine-tuned mathematical functions of *R*_5_ might explain the decrease of *R*_5_ at late times. Yet, there is biological evidence to support that the Rab5 decrease and the Rab7 decrease are not independent events. Possibly, more sophisticated mathematical models, as those explored in Ref. [19], might provide better agreement with the experimental data. Still, simply adding more mathematical terms (and so more parameters) to the dynamical equations would obscure our ability to systematically select between plausible biological mechanisms regulating the dynamics of Rab5 and Rab7.

### A quasi-linear approximation to describe experimental data

Ziegler *et al*. in Ref. [26] considered Rab5-dependent fusion and fission of endosomes, making use of a differential equation similar to Eq. 22. Del Conte-Zerial *et al*. in Ref. [19] modelled the conversion of early endosomes into late endosomes, which assumes the dynamical replacement of Rab5 by Rab7 during endosome maturation. Their equations, summarised in Section 2 of the Supporting information, require a larger number of parameters than those in the cut-off switch model, Eqs. (22)-(24), but do not include a dynamical equation for the number of endosomes. In Fig. 4B we show the fit of their model to the DENV data. Comparison of Fig. 3 and Fig. 4 shows that a model which includes the dynamics of the number of endosomes can explain the molecular mechanisms parsimoniously. Our model does not explicitly consider inactive Rabs ([Rab5-GDP] and [Rab7-GDP]) because their levels remain almost constant (see Ref. [19], Supplementary information). On the other hand, our framework provides dynamical equations for the variance of the number of molecules, as shown in Fig. 4A and Fig. 4C.

In order to perform quantitative model selection, we show in Fig. 4D the value of the Akaike Information Criterion (AIC) for the three models, defined as:

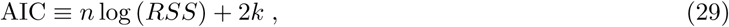

where RSS is the residual sum of squares, *n* the number of points in the data series and *k* the number of parameters in the mathematical model. Similarly, we compute the Bayesian Information Criterion (BIC), defined as

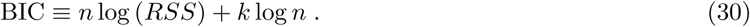

Both methods quantify the goodness of fit but introduce a *penalty* on the number of parameters (the lower the better). As the RSS is similar in all the models (and *n* = 100 is the same for all of them), the most decisive factor is the number of parameters, *k*, thus, pointing at the reduced model, given by Eq. (69) and Eq. (70) (see Fig. 4C). We conclude then that a mathematical that considers the mean number of endosomes in a quasi-steady state, 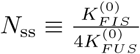, and assumes linear dynamics for the evolution of Rab5 and Rab7 (see Eq. (69) and Eq. (70)) is the best candidate for future model extensions. Clearly, the toggle-switch model cannot capture the behaviour of the experimental data, as argued in the previous section. This emphasises an important conclusion which can be derived from our analysis: fusion, fission and Rab7 inhibition of Rab5 are the main mechanisms regulating endosome maturation and, in our context, the endosome acidification which drives viral escape.

### Fluctuations explain variability of pH-driven viral escape

There exists a simple connection between the levels of endosomal Rab5/Rab7 and endosomal pH given by Eq. (1) (for details, please, see Ref. [14]). In our case, the estimated time evolution of pH is given in Fig. 5A, where we have used the best-fit parameters from Fig. 3 (see Table 1). The shape of the curve is consistent with previous results [14, 25, 41]. Since we can compute the second order moments of the distribution, we can evaluate the standard deviation of the pH (dashed lines in Fig. 5A).

As we mentioned in the Introduction, many intra-cellular processes are triggered by low (below threshold) values of the pH [8–10]. Yet, as shown in Fig. 5A, the pH fluctuates due to the rich dynamics of endosome maturation and Rab decoration. This variability can only be accounted for if one considers the collective dynamics of the population of endosomes, as described by the distribution *n*(*x*_5_, *x*_7_; *t*). We can exploit this variability to define the probability of the pH being below a certain threshold. In particular, we can approximate the fluctuations by a normal distribution, as follows

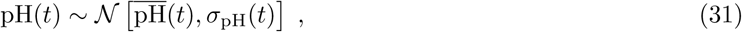

where ∼ 𝒩 denotes normally distributed, and 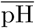 and *σ*_pH_ are the mean and standard deviation of the pH (solid and dashed lines in Fig. 5A, respectively). In Fig. 5B we show synthetically generated histograms according to Eq. (31) to emphasise the role of the width of the distribution and how it affects the probability of having a pH below a certain threshold. The resulting probabilities are shown in Fig. 5C for different values of the pH threshold. For instance, inspection of the green dashed line in Fig. 5C shows that at time 500 seconds, the probability of finding endosomes with pH below 5.5 is already 10%, in spite of the fact that the mean pH is above 6.0 at that time (see the arrows in Fig. 5). This probability represents also the normalised number of viral escape events; that is, for a virus and a fiducial (fixed but arbitrary) pH escape threshold, it gives the probability of viral to escape. For instance, under ideal conditions, a virus that requires a pH=4.5 to escape, would only have a maximal probability of success of 0.022 (∼ 2%). This, in addition to the large mean time to achieve that probability (that would allow the endosome to fuse with the lysosome and, thus, destroy the intra-cellular virus), would make the infection non-viable.

This result is of great relevance in the understanding of endosomal viral escape events. For instance, in Ref. [8] (whose data we are using in the present work), analysis of the experiments revealed that escape events occurred from 300 seconds post-entry (viral endocytosis). Moreover, colocalisation of escape events with levels of Rab5/Rab7 also showed that around 5% of the viral particles escaped from within Rab5-positive early endosomes (lacking Rab7). This implies that the quick pH drop after those early endosomes merged with more acidic endosomes is ultimately regulating the initiation of viral escape events.

## Discussion and conclusions

Traditional mathematical methods, based on ordinary differential equations, can only tell only a part of the story when describing systems with a small number of “particles”, such as endosomes in the present case, and where heterogeneity can be large. Deterministic approaches can be a good approximation if one is interested in average numbers or trends, but exclusively at the cell population level. However, in those cases where the response of a system to small variations of a parameter is abrupt (as in the case of viral escape to a pH threshold), averaging can provide confounding answers or lead to mathematical models with a large number of parameters. One of the main results of the present work is that, while detailed models of the Rab5/Rab7 dynamics can be found to fit accurately to experiments, such as in Refs. [19, 26], a description based on individual endosomes, decorated with Rab molecules (defined by the distribution *n*(*x*_5_, *x*_7_; *t*)), and their interactions (characterised by fusion and fission events), provides a natural link to the underlying biological mechanisms. The mathematical framework proposed here also allows us to characterise the fluctuations beyond mean number of endosomes or Rab molecules, since higher order moments can be computed from the endosomal distribution *n*(*x*_5_, *x*_7_; *t*). As such, we are able to determine the time course of the variance in the number of Rab molecules and the covariance.

We have shown that fusion and fission events regulate the maturation and dynamics of endosomes. Even when we have only considered linear constitutive equations, such as 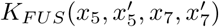 or 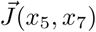, they have allowed us to capture the complexity of the problem at hand. For instance, more detailed models where kiss-and-run [4] or vesicle budding events [42], decoupled from the rates of fusion/fission, might help to quantify the relative role of those mechanisms. Our approach also sheds some light on the variability of other processes relying on this maturation. As an illustration, as it has been shown for many different viruses [8–10], viral escape (after pH drop) is a highly variable process rather than a smooth one, despite the fact that acidification occurs gradually at the endosome level. For instance, in the case of DENV [8], it was already reported that some viruses escaped as early as a few seconds after entry via endocytosis and that, in those cases, fusion with a more acidic endosome preceded that escape event. So, understating endosome dynamics can be relevant to ascertain the role of different entry pathways in the subsequent fate of the virus, since different receptors deliver the virus into distinct populations of early endosomes [43].

Finally, form a practical viewpoint, while Eq. (15) has proven useful to study the time evolution of the number of endosomes, and the total number of active Rab5/Rab7 molecules on the endosomal membrane, it is still a complex system of ODEs, hard to solve analytically. Thus, computational methods aimed to solve these equations can provide rather valuable information. For example, knowledge of the exact distribution *n*(*x*_5_, *x*_7_; *t*) would provide the cell *endosomal pH spectrum*, namely, the number of endosomes with a certain pH, *n*(pH, t), that could be compared with recent experimental methods aimed to quantify intra-cellular pH [37]. Also, this formalism can be translated to other contexts or scales: *n*(*x, y*; *t*) might be seen as the number of cells with a certain expression level of receptors *x* and *y*. Thus, solving the corresponding Smoluchowski equation for *n*(*x, y*; *t*) would be a theoretical metaphor of flow cytometry experiments. Another interesting and timely application of our mathematical framework is that of mitochondrial dynamics and interactions. In this case, and in analogy with the endosomes, mitochondria are subject to fusion and fission events modulated by different cargo species (*e*.*g*., Ca^2+^, ATP, reactive oxygen species, mtDNA, etc.). These extensions will be the aim of future work and out of the scope of the present paper.

## Acknowledgments

This work is partially supported by Ministerio de Economía y Competitividad, Ministerio de Ciencia, Innovación y Universidades, Agencia Estatal de Investigación through Grants FIS2016-78883-C2-2-P and PID2019-106339GB-I00 (MC). The research leading to these results has also received funding from the People Programme (Marie Curie Actions) of the European Union’s Seventh Framework Programme FP7/2007-2013/ under REA grant agreement number 317893 (MC, GL and CMP) and from EPSRC under a University of Leeds Impact Acceleration Account number IAA3025 (GL and CMP).

## Supporting information

### 1 Two-dimensional Laplace transform and moment equations

In analogy with the Laplace transform in one dimension, we introduce the two-dimensional Laplace transform, as follows

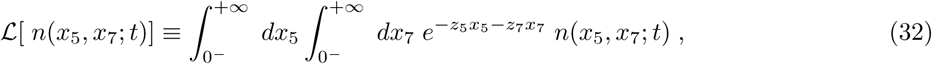

which is a function of the variables *z*_5_ and *z*_7_, and time *t*. We introduce the following notation for the two-dimensional Laplace transform

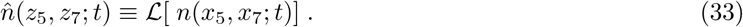

We can also define *partial* (or one-dimensional) Laplace transforms associated with each variable, *x*_5_ and *x*_7_, as follows

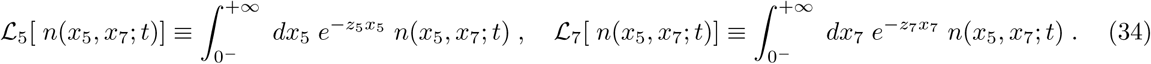

Eq. (32) allows one to derive useful expressions to rewrite the Boltzmann equation, Eq. (15), in terms of the Laplace transform of *n*(*x*_5_, *x*_7_; *t*). Thus, if we make use of the notation introduced in Eq. (33), it can be shown that the following expressions hold [44]:

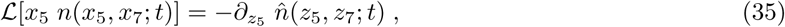

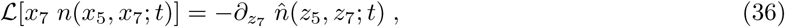

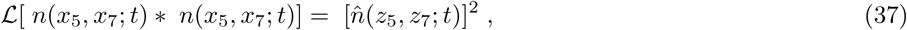

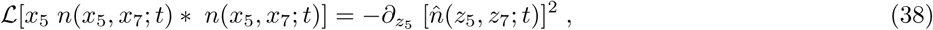

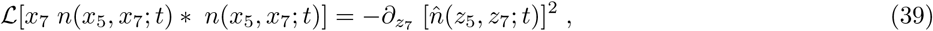

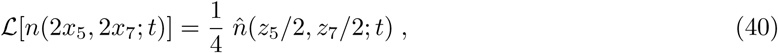

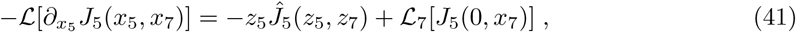

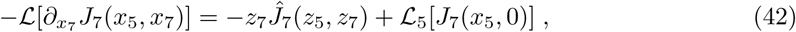

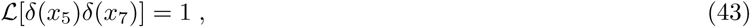

where *Ĵ*_*k*_ (*x*_5_, *x*_7_) is the Laplace transform of *J*_*k*_ (for *k* = 5, 7) and the symbol “*” denotes the convolution; that is,

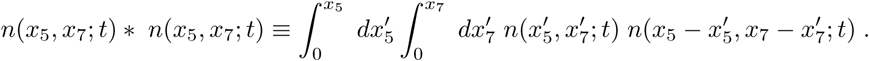

We note that the boundary conditions (see Eq. (14)) imply

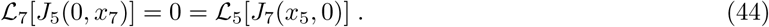

We now make use of the above properties to compute the Laplace transform of Eq. (15). We do so, term by term, as follows:

- **Time derivative:** as the Laplace transform does not involve the time variable, *t*, we have:

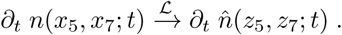
- **Fusion (1):** we make use of Eq. (37),

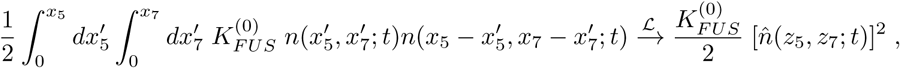

since 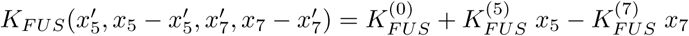.
- **Fusion (2):** we make use of Eq. (38)

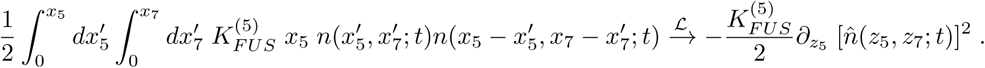
- **Fusion (3):** we make use of Eq. (39)

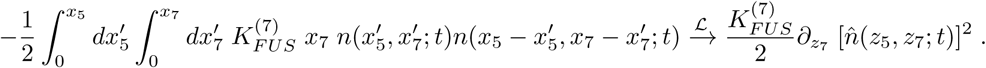
- **Fusion (4):** we make use of the definition of *N* (*t*) in Eq. (16)

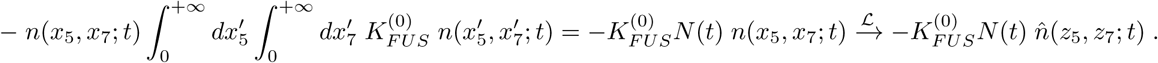
- **Fusion (5):** we make use of Eq. (35) and the definitions of *N* (*t*) and *R*_5_(*t*) in Eq. (16) and Eq. (17), respectively

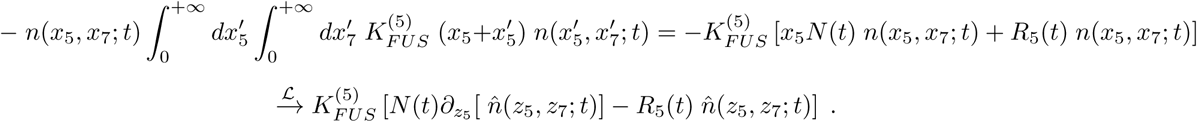
- **Fusion (6):** we make use of Eq. (36) and the definitions of *N* (*t*) and *R*_7_(*t*) in Eq. (16) and Eq. (18), respectively

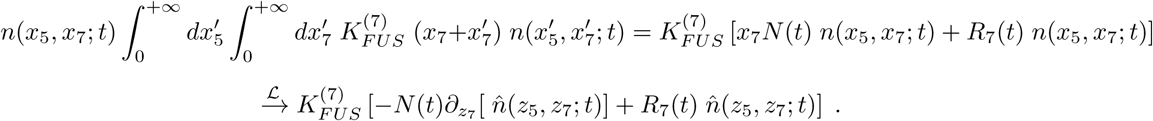
- **Fission (1):** we make use of Eq. (8) and Eq. (40)

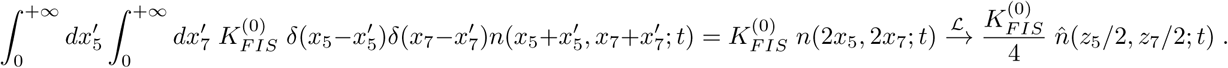
- **Fission (2):** we make use of Eq. (8)

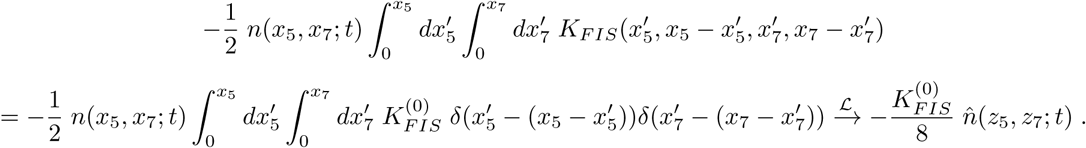
- **Degradation:** we make use of Eq. (15) and Eq. (32)

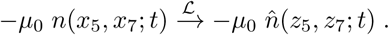
- **Divergence of the current (1):** we make use of Eq. (41) and Eq. (44)

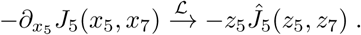
- **Divergence of the current (2):** we make use of Eq. (42) and Eq. (44)

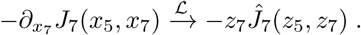
- **Endocytosis:** we make use of Eq. (15) and Eq. (43)

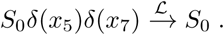

Now that we have established, term by term, the two-dimensional Laplace transform of the Boltmann equation, we take a look at the individual mathematical models considered for Rab5/Rab7 activation/deactivation, the cut-off switch and the toggle-switch models.

#### 1.1 Cut-off switch model

We first consider the cut-off switch model and the precise expression of the divergence of the current under the two-dimensional Laplace transform. We make use of Eq. (10) and Eq. (11) to write

- **Rate of change of Rab5:** from the definition of *J*_5_, Eq. (10), and Eq. (35) and Eq. (36)

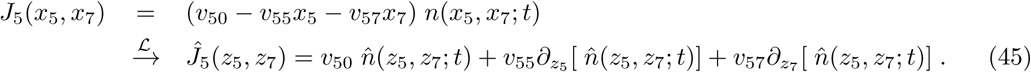
- **Rate of change of Rab7:** from the definition of *J*_7_, Eq. (11), and Eq. (35) and Eq. (36)

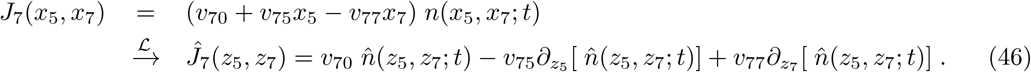

We are now ready to bring all the previous results together for the cutt-off switch model. We find the Laplace transform of Eq. (15) is given by

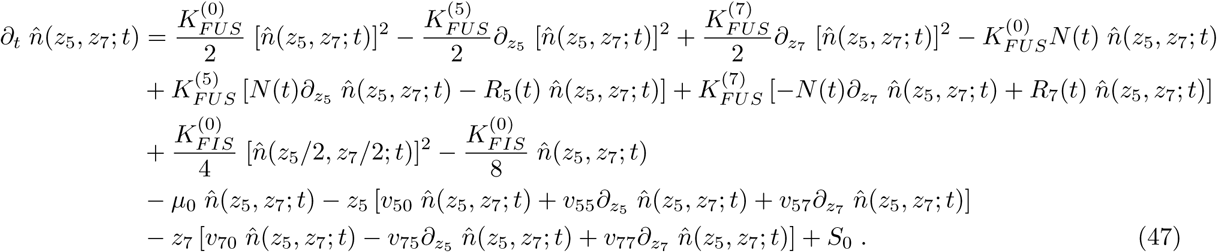

The differential equations for the first order moments, Eq. (22), Eq. (23) and Eq. (24), can be derived from Eq. (47) after one makes the following identifications

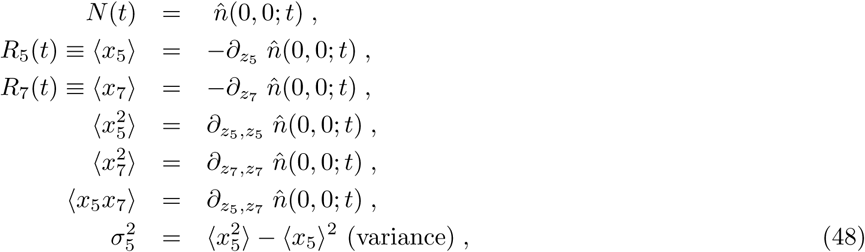

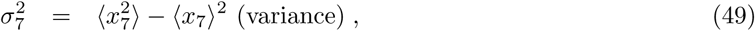

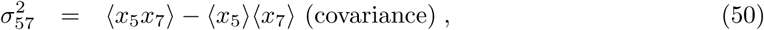

where the derivative has been taken before setting *z*_5_ = 0 = *z*_7_.

The equations for the second order moments, variances and covariance, can be computed in the same way as those for the first order moments, and are given by

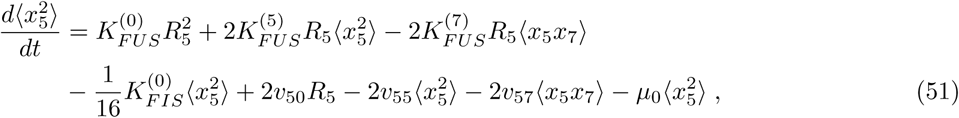

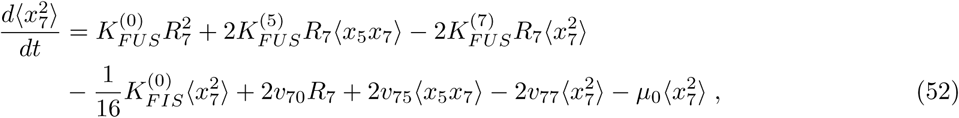

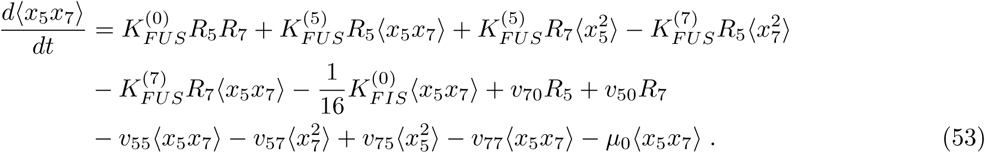

These three ordinary differential equations, together with equations (48)-(50), allow us to obtain the time course of the standard deviations of *R*_5_ and *R*_7_ (and, indirectly, of the endosomal pH).

#### 1.2 Toggle-switch model

In the case of the toggle-switch model, the precise expressions for the current are non-linear. We make use of Eq. (12) and Eq. (13) to write

- **Rate of change of Rab5:** from the definition of *J*_5_, Eq. (12), and Eq. (35) and Eq. (36)

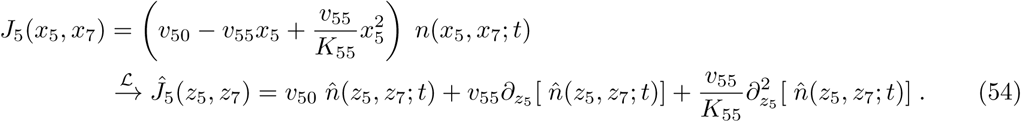
- **Rate of change of Rab7:** from the definition of *J*_7_, Eq. (13), and Eq. (35) and Eq. (36)

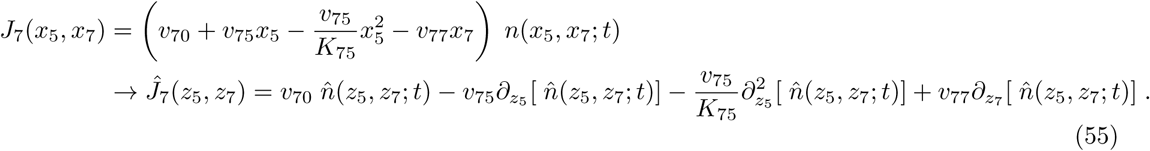

We are now ready to bring all the previous results together for the toggle-switch model. We find the Laplace transform of Eq. (15) is given by

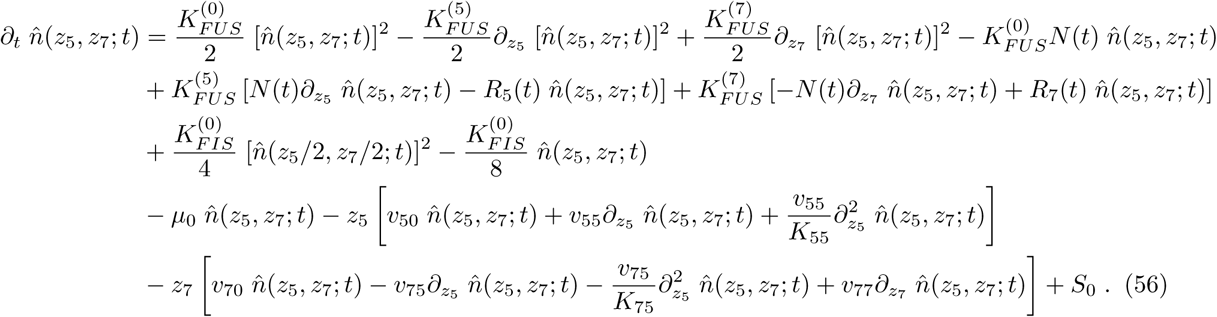

The differential equations for the first order moments, Eq. (25), Eq. (26) and Eq. (27), can be derived from Eq. (47) in the same way as for the cut-off switch model. We note that the differential equation for the mean number of endosomes, *N* (*t*), is the same for both models, since the specific form of the currents, *J*_5_(*x*_5_, *x*_7_) and *J*_7_(*x*_5_, *x*_7_), does not change the mean number of endosomes. However, as it should be expected, the differential equations for *R*_5_(*t*) and *R*_7_(*t*) depend on the choice of currents. We have

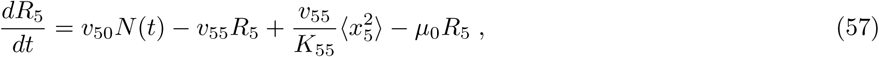

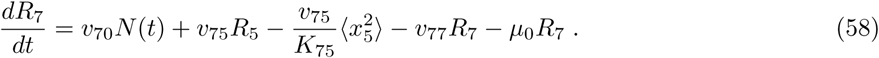

We note that the non-linear nature of the currents in this model implies that the differential equations for the first order moments, Eq. (57) and Eq. (58), depend on the second order moments. At the level of the Laplace transform, this non-linearity implies that (for *k* = 5, 7) *Ĵ*_*k*_ involves second order derivatives of 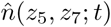. Thus, the equations for the first order moments involve the second order ones, and so on. If we define the joint cumulants [45], *κ*_*i,j,k*_, where *i, j, k* = 5, 7, as follows:

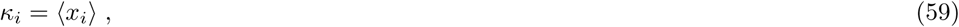

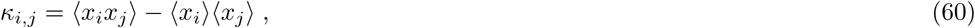

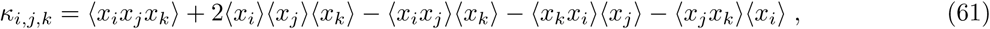

then we can, for the sake of simplicity, make use of a zero-cumulant moment-closure approximation [46]. This approximation implies the following choices for the relevant cumulants

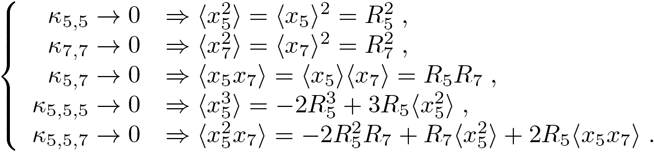

If we make use of the zero-cumulant moment-closure approximation above in Eq. (57) and Eq. (58), we obtain Eq. (26) and Eq. (27), respectively. Similarly, for the second order moments we find the following equations

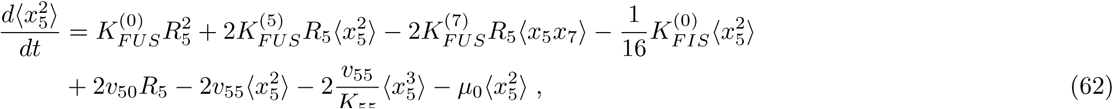

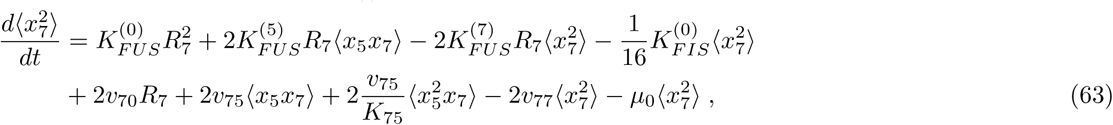

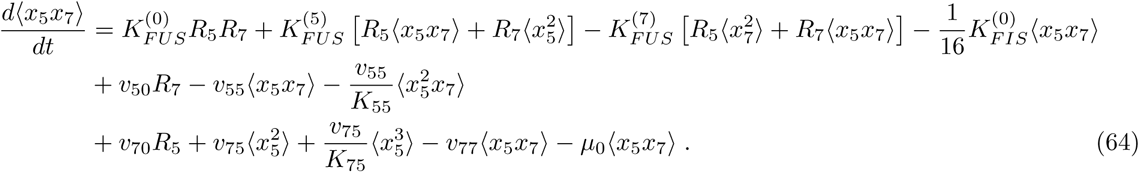

If we now make use of the moment-closure approximation, the previous equations can be written as follows:

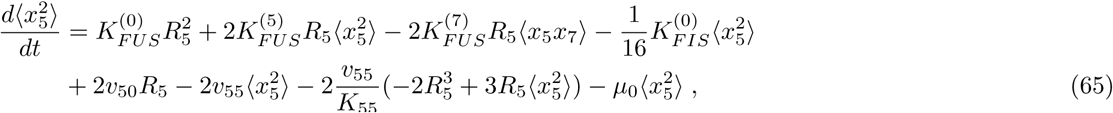

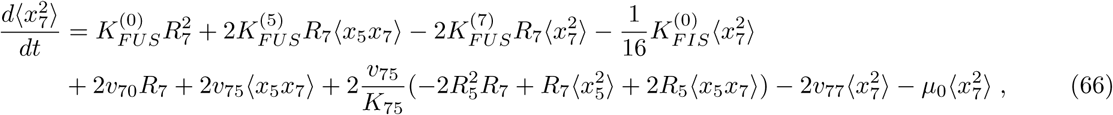

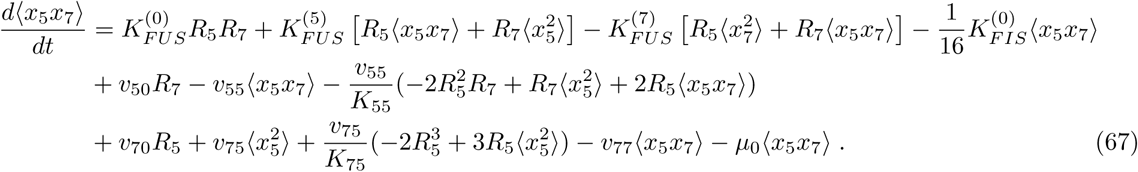

### 2 Fitted parameters and sensitivity analysis

We have made use of the software Copasi to perform parameter fitting with three different methods: Levenberg-Marquardt, steepest descent, and Hooke and Jeeves methods [47]. The results converged in all three cases. In Table 1, we summarise the best-fit parameters for the three models introduced in this paper.

We also used the Copasi built-in (relative) sensitivity analysis algorithm. Tables 2-5 provide a summary of the results of that algorithm for the three models used: cut-off switch model, toggle-switch model and the model in Ref. [26]. In each table, the column *Aggregated* is the square root of the sum of the squares of each sensitivity. This measure (also computed by Copasi) gives an idea of the most relevant parameters (highest value in that column). Mathematically, we have

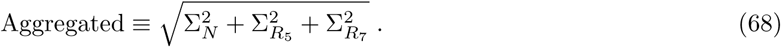

Consider, for instance, the variable *N* and the parameter *S*_0_. The value showed in the table corresponds to

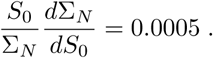

That is, every order of magnitude that we increase *S*_0_ only produces an increase of 10^0.0005^ ≃ 1.001 in the mean number of endosomes (a mere ∼ 0.1% increase). Thus, values close to 1 correspond, roughly, to a linear dependence between variable and parameter and values close to −1, an inverse proportionality. In addition, we have added the column *Normalised*, where we divide the *Aggregated* column by the maximum of all parameters. For instance, in Table 2, the maximum is 1.7294 corresponding to 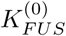.

From Table 2 we can derive a simpler model, where we drop the less sensitive parameters, namely, *S*_0_, *v*_70_ 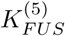 and 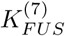. We have also set the number of endosomes to a constant, given by the steady state value of the full model, and as given by Eqs. (22)-(24). The resulting model is given by the following equations

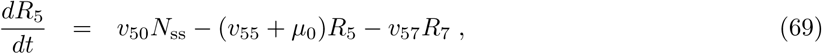

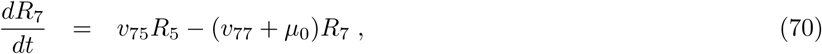

with parameters shown in Table 1 and sensitivities in Table 3. Note that, from Table 2, only 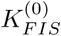 and 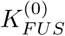 have strong sensitivities on the variable *N*. We can then safely assume that

**Table 3.** Sensitivity analysis of the reduced model computed by three different methods implemented in the software Copasi [47]. The column *Aggregated* is defined in Eq. 68. The column *Normalised* is the value in *Aggregated* divided by the maximum value in that column.

| Parameter | $\Sigma_N$ | $\Sigma_{R_5}$ | $\Sigma_{R_7}$ | Aggregated | Normalised |
| --- | --- | --- | --- | --- | --- |
| $K_{FIS}^{(0)}$ | 1.0000 | 0.9995 | 1.0001 | 1.7318 | 1.0000 |
| $K_{FUS}^{(0)}$ | -0.9990 | -0.9992 | -0.9991 | 1.7305 | 0.9992 |
| $v_{50}$ | $< 10^{-4}$ | 0.9991 | 1.0013 | 1.4145 | 0.8168 |
| $v_{57}$ | $< 10^{-4}$ | -0.7805 | -0.7288 | 1.0679 | 0.6166 |
| $v_{75}$ | $< 10^{-4}$ | -0.7805 | 0.2704 | 0.8260 | 0.4770 |
| $v_{77}$ | $< 10^{-4}$ | 0.7556 | -0.2928 | 0.8103 | 0.4679 |
| $v_{55}$ | $< 10^{-4}$ | -0.1181 | -0.3022 | 0.3245 | 0.1874 |

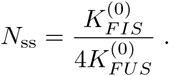

This is confirmed in Fig. 6, where the number of endosomes quickly converges to its steady state. Finally, and for the sake of completeness, in Table 4 we summarise the sensitivity analysis for the toggle-switch model.

**Table 4.**
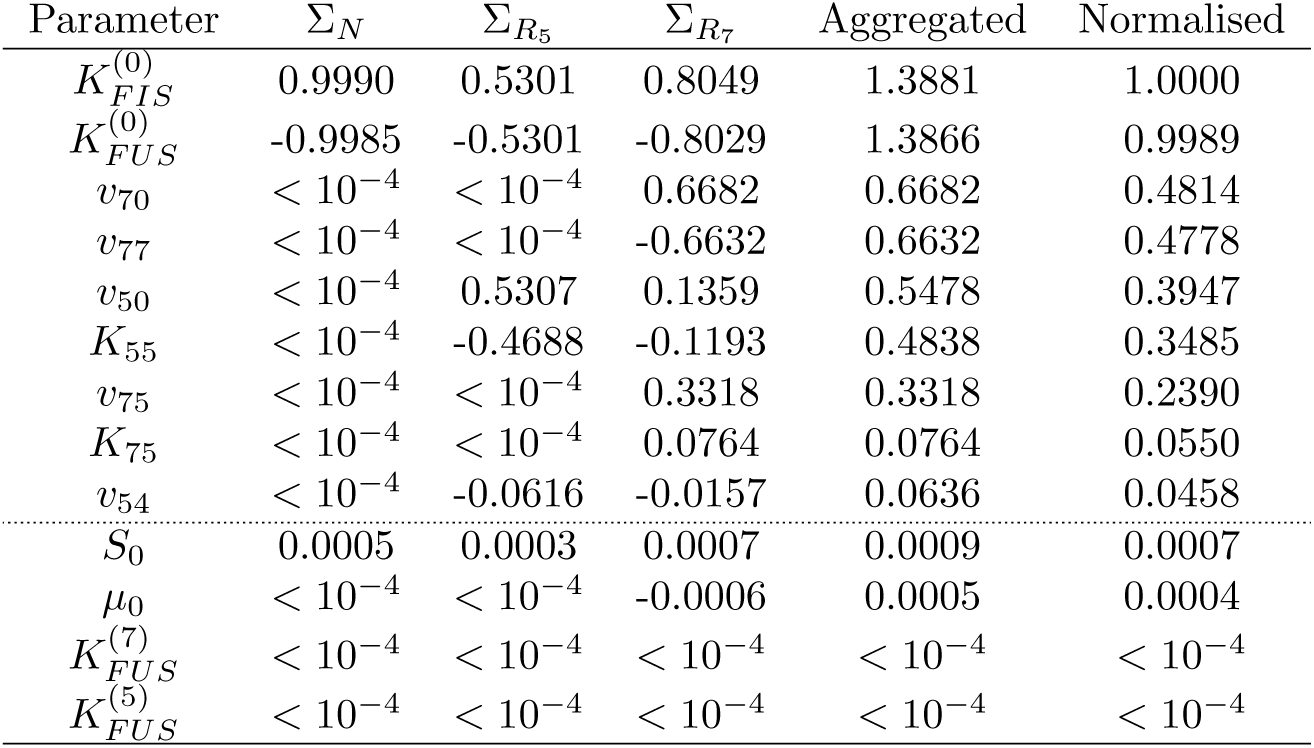
Sensitivity analysis for the toggle-switch model computed by three different methods implemented in the software Copasi [47]. The column *Aggregated* is defined in Eq. 68. The column *Normalised* is the value in *Aggregated* divided by the maximum value in that column. The thin dotted lines are a guide to the eye to separate the most sensitive parameters (top part of the table) and the least (bottom part).

**Table 5.** Sensitivity analysis of the model in Ref. [19] (corresponding to Eqs. (71)-(74) of Section 2 in Supporting information) obtained by three different methods implemented in the software Copasi [47]. The column *Aggregated* is defined in Eq. 68. The column *Normalised* is the value in *Aggregated* divided by the maximum value in that column. The thin dotted lines are a guide to the eye to separate the most sensitive parameters (top part of the table) and the least (bottom part).

| Parameter | $\Sigma_{Rab5-GDP}$ | $\Sigma_{Rab5-GTP}$ | $\Sigma_{Rab7-GDP}$ | $\Sigma_{Rab7-GTP}$ | Aggregated | Normalised |
| --- | --- | --- | --- | --- | --- | --- |
| $k_7$ | 0.0022 | 28.8007 | -1.0044 | -7.8522 | 29.8688 | 1.0000 |
| $k_{1,7}$ | 0.0021 | 28.7917 | -0.0055 | -7.8498 | 29.8426 | 0.9991 |
| $v_7$ | -0.0018 | -27.1203 | 1.0046 | 7.57364 | 28.17588 | 0.9433 |
| $k_{e,7}$ | -0.0018 | -26.6033 | 0.0045 | 7.4276 | 27.6207 | 0.9247 |
| $k_{g,7}$ | 0.0007 | 9.4375 | -0.0020 | -2.5938 | 9.7875 | 0.3277 |
| $k_{g,57}$ | 0.0004 | 4.3526 | -0.0010 | -1.1988 | 4.5147 | 0.1512 |
| $k_{f,57}$ | 0.0003 | 3.3470 | -0.0007 | -0.9223 | 3.4718 | 0.1162 |
| $k_{g,75}$ | -0.0005 | 0.2003 | 0.0012 | 1.6523 | 1.6643 | 0.0557 |
| $h_7$ | 0.0003 | 1.2028 | -0.0007 | -0.3313 | 1.2476 | 0.0418 |
| $k_5$ | 0.9998 | 0.0216 | 0.0002 | 0.2905 | 1.0414 | 0.0349 |
| $v_5$ | -0.9989 | -0.0135 | -0.0003 | -0.2928 | 1.0410 | 0.0349 |
| $k_{f,75}$ | -0.0002 | 0.0541 | 0.0006 | 0.6927 | 0.6948 | 0.0233 |
| $k_{e,57}$ | 0.0001 | -0.5757 | 0.0001 | 0.1589 | 0.5972 | 0.0200 |
| $k_{e,5}$ | -0.0002 | 0.0205 | 0.0002 | 0.2908 | 0.2915 | 0.0098 |
| $k_{e,75}$ | 0.0001 | -0.0133 | -0.0002 | -0.2268 | 0.2272 | 0.0076 |
| $k_{g,5}$ | $< 10^{-4}$ | -0.0074 | -0.0001 | -0.1559 | 0.1561 | 0.0052 |
| $k_{1,5}$ | $< 10^{-4}$ | -0.0068 | -0.0001 | -0.1354 | 0.1355 | 0.0045 |
| $k_{f,5}$ | $< 10^{-4}$ | -0.0007 | $< 10^{-4}$ | -0.0661 | 0.0661 | 0.0022 |
| $T_0$ | $< 10^{-4}$ | 0.0053 | $< 10^{-4}$ | -0.0390 | 0.03938 | 0.0013 |

### 3 Study of two previous of Rab dynamics

#### 3.1 Mathematical model in Ref. [19]

The authors in Ref. [19] encoded the dynamics of Fig. 1 as a set of four ODEs, where each reaction term was fitted to different mathematical equations. One set of functions ^3^ provides the following equations (with *t* the variable describing experimental time)

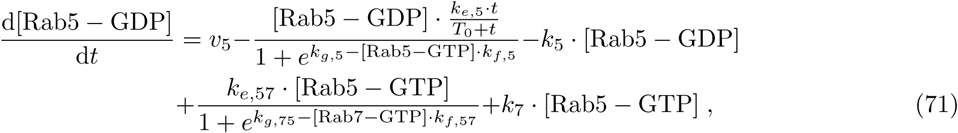

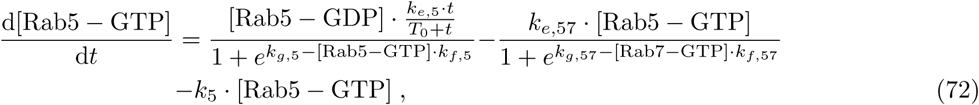

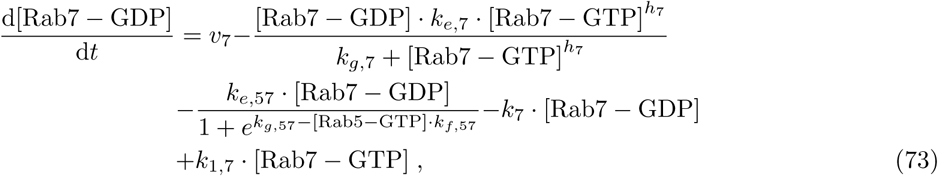

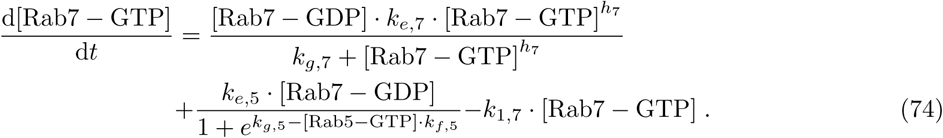

Although this model looks clearly more complex than our model (see Eq. (23) and Eq. (24)), numerical integration of these equations (as shown in Fig. 4B) shows that the variables [Rab5-GDP] and [Rab7-GDP] are almost constant (not shown). Hence, Eqs. (71)-(74) are identical to Eq. (23) and Eq. (24), after linearisation of the equations for [Rab5-GTP] and [Rab7-GTP], with the following approximations:

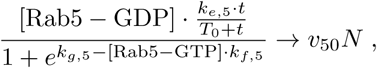

and

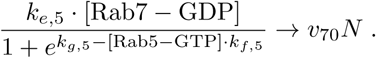

#### 3.2 Mathematical model in Ref. [26]

For completeness, we reproduce here the system of equations in Ref. [26]. Note that the first two terms in Eq. (22) are equivalent to those in Eq. (78).

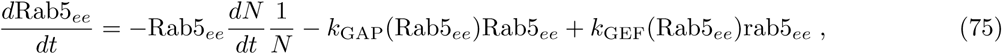

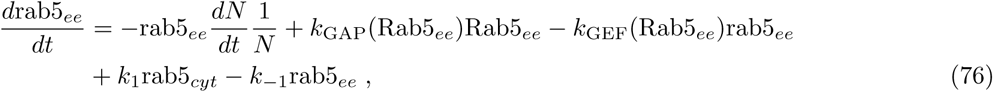

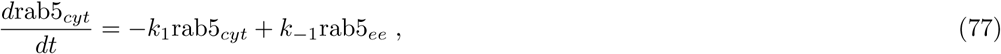

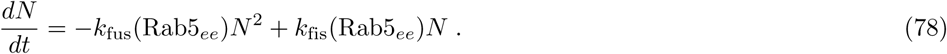

Note that, as according to our model, the number of endosomes reaches quickly a steady state, the terms proportional to 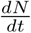 are negligible in comparison to the other terms. Similarly, as shown in the Supporting information in Ref. [19], the inactive *rab*5_*ee*_ is also almost constant. Thus, we can reduce the equations above to the following system

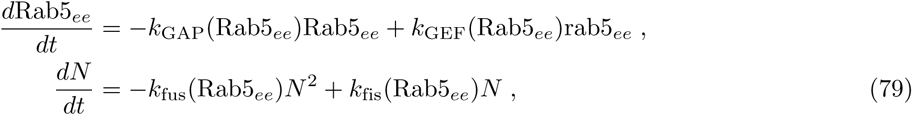

where our notation *k*_GAP_(Rab5_*ee*_), *k*_GEF_(Rab5_*ee*_), *k*_fus_(Rab5_*ee*_) and *k*_fis_(Rab5_*ee*_) implies that these rates are functions of the variable Rab5_*ee*_. Our analysis has shown that the source term, proportional to *S*_0_, and the death term, proportional to *µ*_0_, are only important in the initial and transient regime. So Eq. (23) and Eq. (22) can be written as:

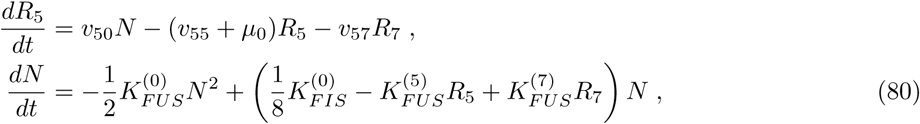

which are analogous to Eqs. (79). In our case, the inclusion of the variable *R*_7_ (not included in Ref. [26]) can explain the need to choose more sophisticated mathematical functions for *k*_*GEF*_ and *k*_*GAP*_ in the equations above. To illustrate this point, in Fig. 7 we show Rab5 as a function of Rab7 for both the experimental data (circles) and the fitted model (solid line). This allows us to conclude that Rab5 depends non-linearly on Rab7. If we describe Rab5, without reference to the dynamics of Rab7, as done in Ref. [26], one would require an equation for *R*_5_ containing non-linearities. In particular, if we denote by *k*_*GAP*_ (*Rab*5_*ee*_), the coefficient of *Rab*5_*ee*_ and by *k*_*GEF*_ (*Rab*5_*ee*_), the coefficient of *rab*5_*ee*_, one can see that the dependence on *R*_7_ in the first case and on *N* on the second one, are equivalent to the non-linear functions of *R*_5_ in Eq. (79).

**Fig 7.**
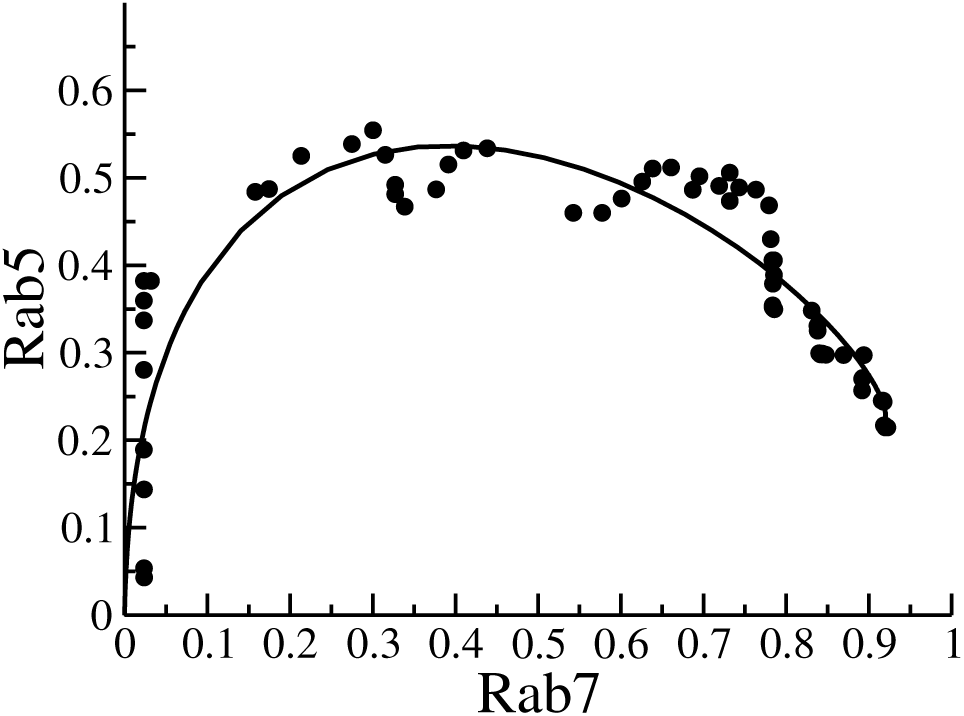
Phase-diagram of Rab5 and Rab7. Circles: experimental data. Solid line: fit to Eq. (22) and Eq. (24). This shows that *R*_5_ can be expressed as a non-linear function of *R*_7_. Thus, linear terms of *R*_5_ and *R*_7_ can be misidentified with non-linear functions of *R*_5_ alone.

Finally, and as shown in the Supplementary Figures 1a-1b from Ref. [19], the best-fit for the fusion rate is almost independent of Rab5 (note the large bars in logarithmic scale), while the best-fit for the fission rate consistently changes as a function Rab5. In our framework, this dependence is encapsulated in the factor 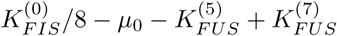 multiplying the variable *N* in Eq. (22), and the constant coefficient 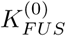 multiplying *N* ^2^. So the we can identify

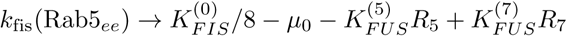

in Eq. 79. This identification clearly shows that *k*_fis_(Rab5_*ee*_) is a non-linear function of *R*_5_ since *R*_7_ = *R*_7_(*R*_5_) (see Fig. 7). This completes the connection between the mathematical framework presented here and previous mathematical models of endosome maturation.

## Footnotes

1 In order to avoid confusion, in this manuscript we shall refer to viral RNA release from the endosome to the cytosol as *escape* and to the merging of two endosomes as *fusion*. We note that other authors refer to fusion as the merging of the viral envelope and the endosomal membrane.

2 We note that *v*_57_ = 0 in the toggle-switch hypothesis, since Rab7 does not affect the dynamics of Rab5.

3 As documented in http://biomodels.caltech.edu/BIOMD0000000174.

